# A contractile injection system mediates developmentally regulated intra-strain cell death in *Streptomyces coelicolor*

**DOI:** 10.1101/2022.08.10.503483

**Authors:** Maria Vladimirov, Ruo Zhang, Stefanie Mak, Justin R. Nodwell, Alan R. Davidson

## Abstract

Extracellular contractile injection systems (eCIS) are encoded in diverse clades of bacterial species. Although closely related to contractile phage tails, these entities can inject toxic proteins into eukaryotic cells. The roles of eCIS in mediating cytotoxic activities has led to a view of them as defense mechanisms that are not central to normal bacterial lifecycles. Here, we provide evidence that eCIS play an entirely distinct role in *Streptomyces coelicolor* (*Sco*), where they appear to participate in the complex developmental process of this species. In particular, we have shown that *Sco* produces eCIS particles as a part of its normal growth cycle and that strains lacking functional eCIS particles exhibit pronounced alterations in their developmental program. Most intriguingly, eCIS-deficient mutants display significantly reduced levels of cell death and altered morphology during liquid growth. These data suggest that *Sco* eCIS function by inducing intra-strain lethality rather than by attacking foreign species.

## Introduction

Many strains of bacteria encode extracellular contractile injection systems (eCIS)^1–3^. eCIS are closely related to contractile-tailed bacteriophage (phage) tails. Like tails, they are composed of a long tube attached to a baseplate (Fig. 1f). The proteins comprising the eCIS tail-like structure are similar in sequence to phage proteins. However, genomic regions encoding eCIS are distinguished from phage tail regions by invariably encoding an AAA+ ATPase of unknown function and a tail terminator protein (Tr) that is a member of the *Pvc16_N* (PF14065) Pfam family^4,5^. They also often encode recognizable toxin proteins^2,6,7^. All of the eCIS with characterized functions mediate interactions between bacteria and eukaryotic cells. *Serratia* and *Photorhabdus* species release eCIS structures, known as Anti-feeding prophages (Afp) and Photorhabdus Virulence Cassettes (PVC), respectively, that mediate killing of insect cells^8,9^. This killing occurs through the injection of insecticidal toxins encoded downstream of the eCIS operon^8–10^. MACs (Metamorphosis Associated Contractile structures) are a distinct eCIS produced by the marine bacterium, *Pseudoalteromonas luteoviolacea*, that are required for the morphogenesis of the tubeworm *Hydroides elegans* in a mutualistic interaction^11^. The structures of several other eCIS have been examined in detail^12–14^, but their functions have not been determined.

**Fig. 1.**
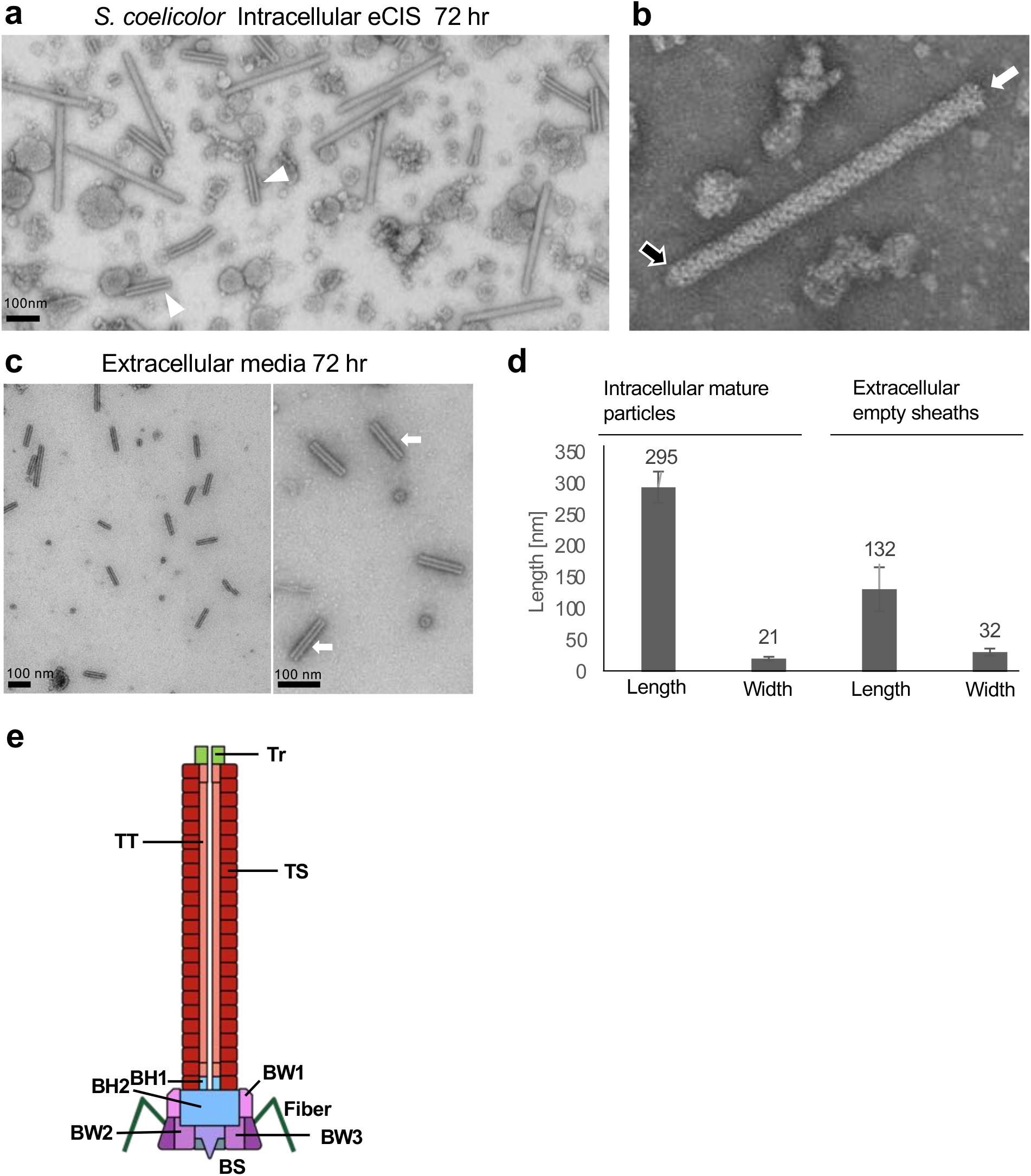
eCIS-derived particles can be visualized in cell lysates and extracellular media. ***a-c***, Transmission electron micrographs represent wild-type *Sco* grown in liquid YEME media for 72 hr post inoculation. **a**, eCIS were purified from *Sco* lysates as described in Methods and concentrated to 30X the concentration of the original culture. Images were taken at 100,000 X magnification. White arrow heads point to empty sheaths, present alongside with fully assembled eCIS. **b**, A single assembled eCIS particle in its uncontracted conformation is shown. The white arrow points to the baseplate, and the black arrow shows the tail terminator complex. **c**, Extracellular fraction. Cell-free media was ultracentrifuged and concentrated to 30X the concentration of the original sample Images taken at 80,000X magnification. White arrows point to typical emptied contracted tail sheath particles. Scale bar represents 100 nm in all images. **d**, Representative TEM images of *Sco* eCIS-derived structures from early stages of liquid growth are shown. All images are of purified lysate samples of wild-type *Sco* grown in liquid YEME media for 48 hr post inoculation. Tails of different aberrant forms are shown. White arrows point to abnormal contraction or breaks in the sheath coating and black arrows point to the exposed internal tail tube. Images were taken at 100,000X magnification. Scale bar represents 100 nm. **e**, Average length and width of the eCIS-derived particles described in (a-c) (n=80 for fully assembled uncontracted intracellular particles, n=215 for extracellular contracted empty sheath particles). Error bars represent standard deviation of the mean. **f**, Schematic illustration of an eCIS particle. The diagram shows the conserved core structural components of an eCIS tail. Proteins are indicated by abbreviations (Tr = tail terminator; TT = tail tube; TS = tail sheath; BH1, BH2 = baseplate hub components; BW1, BW2, BW3 = baseplate wedge components; BS = tail spike).

eCIS regions have clearly been disseminated through horizontal gene transfer and this has resulted in their appearing sporadically across diverse bacterial clades^2,3^. A striking exception to this pattern is seen in the *Streptomyces* genus where the majority of species (>70%) encode at least one eCIS. *Streptomyces* propagate through a complex developmental program. Germinated spores initially grow to form extensive vegetative hyphae by a combination of tip extension and branching^15^. Upon nutrient deprivation or stress, programmed developmental and morphological changes result in formation of aerial hyphae and subsequent formation of spores. This “metabolic switch” is accompanied by a large shift in gene expression and the onset of secondary metabolite production, many of which are antibiotics^16,17^.

Given the uniquely complex lifestyle of *Streptomyces* and the prevalence of eCIS encoding regions in this genus, we hypothesized that eCIS may play a fundamental role in the developmental program of these bacteria. To test this idea, we initiated studies on the eCIS encoding region of the model strain *Streptomyces coelicolor* A3(2) (*Sco*), which has a single eCIS-encoding region. The goals of these experiments were to determine whether eCIS are produced as part of the normal *Sco* lifecycle and what role these little characterized entities might be playing.

## Results

### eCIS tail-like structures can be purified from *Sco* lysates

To determine whether wild-type (WT) *Sco* produces eCIS particles, cell lysates of *Sco* cultures grown in liquid medium for 72 hr were subjected to a standard eCIS purification protocol (see Methods). Examination of the resulting purified samples using transmission electron microscopy (TEM) revealed abundant phage tail-like particles with average lengths of 300 nm and widths of 21 nm (Fig. 1a-b, 1d). These particles closely resemble previously observed eCIS particles with a rounded cap on one end and a baseplate on the other as indicated by arrows (Fig. 1b)^5,18,19^. The baseplate is somewhat smaller than other eCIS and fibers, which are attached to the baseplates of some eCIS^5,11,20^, were not detectable. The purified samples also contained many empty sheath structures of varying length (Fig. 1a, white arrowheads).

Strikingly, TEM micrographs of the extracellular fraction (supernatant collected after cell pelleting) revealed only what appeared to be emptied out sheaths left behind after eCIS contraction. These structures were 100-150 nm long and 28-35 nm wide with a typical striation pattern characteristic of polymerized tail sheath, but they had no density in the center and were wider than mature wild-type eCIS tails (Fig. 1c,d). Tail sheath (TS) protein was also detected in the extracellular (E) fraction by Western blots (Supplementary Fig. 1a,b). The strictly cytoplasmic protein, ActR, was not detected in the extracellular fraction (Supplementary Fig. 1b), suggesting that the appearance of TS protein in this fraction was not due to generalized cell lysis. *Sco* strains bearing deletions in the genes encoding either the tail sheath (ΔTS) or the baseplate wedge proteins (ΔBP) (Fig. 1e) displayed no tail-like structures in extracellular samples or cellular lysates. Western blots confirmed the absence of these proteins from the mutant strain preparations (Supplementary Fig. 1 a,c).

The presence of a gene encoding an AAA+ ATPase is a universal feature of eCIS genomic regions, yet no function had been ascribed to this enzyme. To investigate this issue, we generated a deletion in the *Sco* ATPase gene (*sco4259*) and performed TEM experiments as described above on lysates of this mutant. The eCIS particles produced appeared identical to those produced by the WT strain (Supplementary Fig. 2a). Similar levels of extracellular empty sheaths were also observed (Supplementary Fig. 2b,c). These results imply that the ATPase is not required for the production of eCIS-like particles with normal appearance. This finding agrees with work on the *Photorhabdus* eCIS^5^, which also showed that normal looking eCIS particles are formed in the absence of the ATPase^5^.

We used mass spectrometry to determine the composition the purified tail-like particles produced by *Sco*. As expected, most of the eCIS structural proteins (Fig. 1e) that have phage tail homologues were detected in all three purifications (TR, TS, TT, BH2, BW2) while others of these conserved structural proteins (BH1 and BW1) were detected in at least one of three separate purifications performed (Supplementary Tables 1 and 2). BW3 was the only conserved structural protein that was not detected in any purifications. Two proteins of unknown function, encoded by genes *sco4242* and *sco4256*, were detected in all three purifications, and one other, encoded by *sco4251*, was detected in at least one purification. The AAA+ ATPase was not detected in purified eCIS particles. We also performed mass spectrometry on eCIS-related particles purified from the extracellular fraction. In this case we detected only TS and TT proteins, which are the likely components of the empty contracted sheaths observed by TEM. Importantly, purified preparations produced from the ΔBP mutant contained only TT and TS proteins, demonstrating that the detection of most eCIS proteins requires the proper eCIS assembly, which cannot occur in the absence of the baseplate wedge proteins^5^. Aggregates of TS and TT protein may still be purified by high-speed centrifugation. Overall, these mass spectrometry experiments show that the tail-like particles that we have purified are indeed produced from the eCIS region of *Sco* and that the composition of these particles is the same as other eCIS.

### eCIS do not contribute to killing of other species by *Sco*

A previous study implicated the eCIS of *Streptomyces lividans* as being responsible for a growth inhibitory activity against the yeast, *Saccharomyces cerevisiae*^21^. Since the components of *S. lividans* eCIS are very similar in sequence to those of the *Sco* system, it was of interest to determine whether the *Sco* eCIS also displayed this activity. As shown in Fig. 2, we found that WT *Sco* did indeed inhibit the growth of *S. cerevisiae* and that this activity was not seen for the ΔTS strain. As Sco is known to produce several antibiotics that might contribute to the observed growth inhibition, we tested the ability of *Sco* strain M1152^22^, in which genomic regions required for synthesis of the four major *Sco* antibiotics are deleted (*Δact, Δred, Δcda, Δcpk*), to inhibit yeast growth. We confirmed that this strain still produces eCIS-related particles (Supplementary Fig. 3a). We found that neither strain M1152 nor a mutant version of this strain that does not produce eCIS (M1152ΔTS) inhibited yeast growth (Fig. 2). These results imply that antibiotics are required for the growth inhibition of *S. cerevisiae* and that the absence of eCIS may indirectly affect the inhibitory activity by perturbing antibiotic production.

**Fig. 2.**
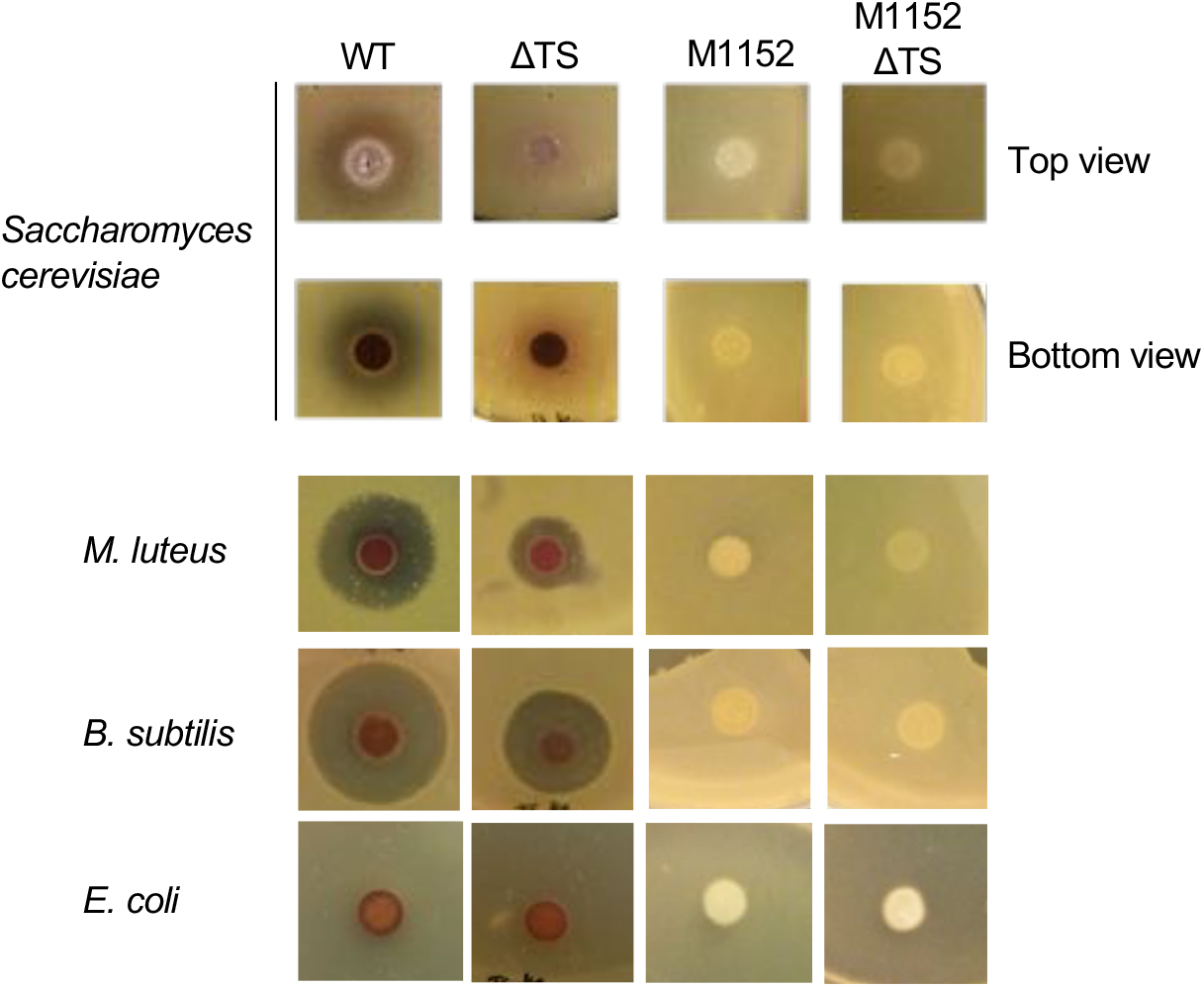
Interspecies growth inhibition induced by WT and eCIS-deficient *Sco* strains. Assays were performed by spotting on agar plates 1×10^5^ SFU of *Sco* wild-type (WT), tail sheath knockout (ΔTS), M1152 (a mutant that does not produce the four major *Sco* antibiotics), or M1152 harboring a tail sheath knockout mutation (M1152 ΔTS). Cells were grown for 3 days before being overlaid with species to be tested as indicated in each panel. Zones of clearing around the *Sco* colonies indicate lethality or growth inhibitory activity against the indicator lawn. All of the results shown for a tested species strain were obtained from the same agar plate. Images are representative examples of many replicates (n > 3).

Similar to our results with *S. cerevisiae*, we found that *Sco* inhibited growth of the Gram-positive bacterial species, *Bacillus subtilis*, and *Micrococcus luteus*, and that this inhibition was partially attenuated in the ΔTS strain (Fig. 2). However, strain M1152 and the ΔTS mutant lacked this activity. Finally, we found that *Sco* inhibited the growth of four different *Streptomyces* species, but the ΔTS strain displayed the same level of inhibition, implying that the eCIS was not required for the growth inhibition (Supplementary Fig. 3b). In summary, these growth inhibition experiments demonstrate that *Sco* does inhibit the growth of some bacterial species and *S. cerevisiae*, but there does not appear to be a direct role for the eCIS in this activity.

### eCIS mutants exhibit decreased hyphal death and reduced pellet size during differentiation in liquid cultures

Given the apparent lack of killing of other species by the *Sco* eCIS, we postulated that eCIS might mediate an intra-strain cell killing activity during the normal developmental program. To visualize and quantify levels of cell death during *Sco* differentiation, we used a well-established bacterial viability assay^23^. Samples are stained with SYTO 9, which labels both live and dead cells, and with propidium iodide (PI), which penetrates and stains only bacteria with damaged membranes, which are likely dead. In the presence of both stains, live bacteria appear green whereas dead bacteria appear red. After 48 hr of growth in liquid media, WT *Sco* grew mycelial pellets as has been previously observed^24,25^. The average diameter of these pellets was 85 *μ*m (Fig. 3a). The same size pellets were seen in the eCIS-deficient strains. By quantitating the staining by SYTO9 and PI, we calculated the ratio of live to dead cells in the WT cultures to be 0.8 (Fig. 3a). The live/dead ratios were slightly less in the mutant cultures though the difference was not significant. Differences between the WT and the eCIS-deficient strains became evident at the 54 and 74 hr of growth points. WT samples developed many large mycelial pellets ranging from 100 to 300 *μ*m in diameter that displayed increased PI staining in the center (Fig. 3b,c). By contrast, in both eCIS-deficient mutants, the preponderance of observed hyphal pellets remained small with the average pellet sizes being half that of WT *Sco*. In addition, the live/dead cell ratios of the mutants were double those of WT (Fig. 3b,c). Overall, these data show that the development of WT *Sco* in liquid media is accompanied by formation of large mycelial pellets and considerable levels of cell death. Mutants lacking eCIS particles significantly deviate from this behavior displaying smaller cell pellets and a much lower frequency of cell death.

**Fig. 3.**
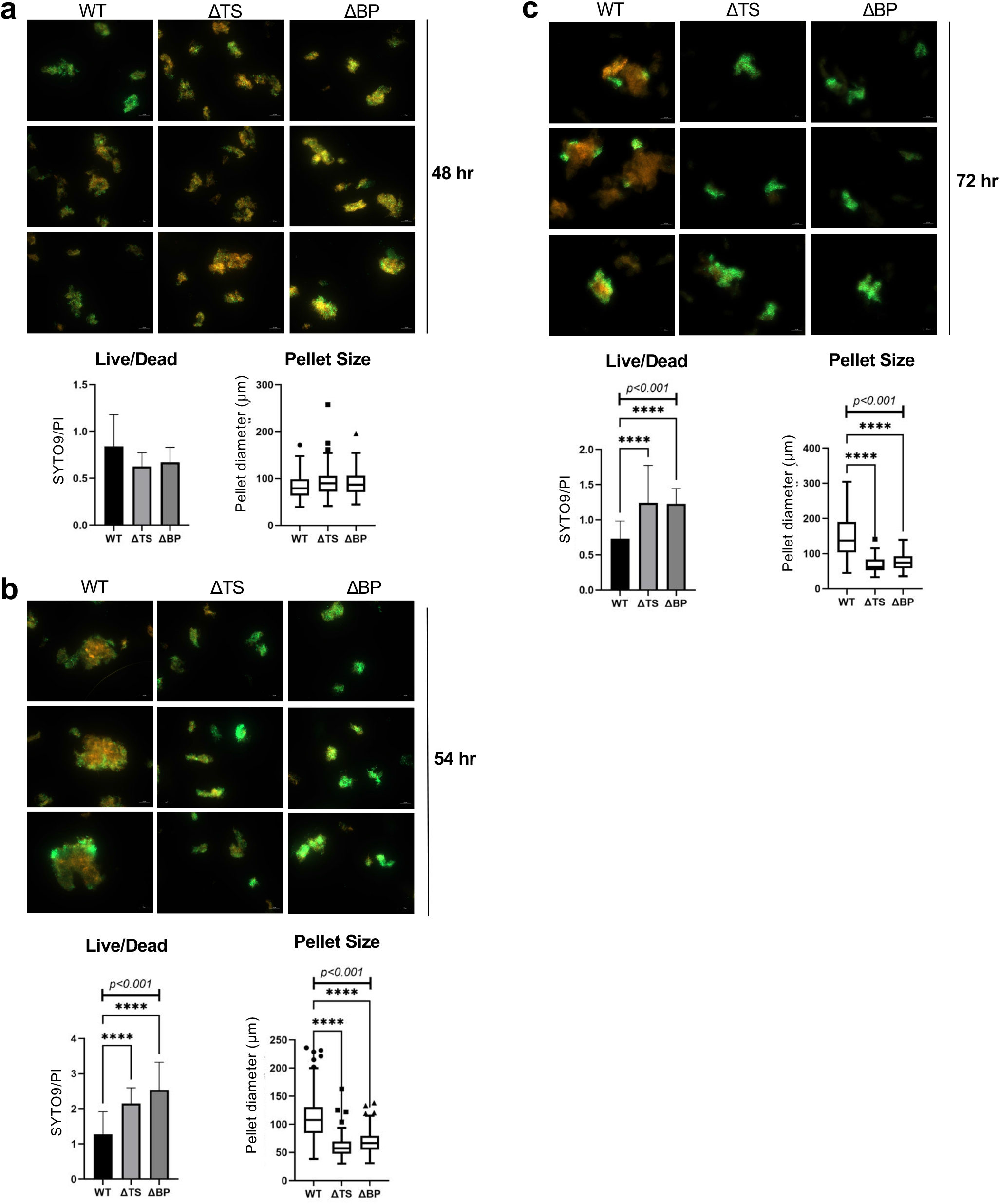
*Sco* eCIS mutants exhibit reduced cell death during hyphal differentiation. *Sco* wild-type (WT), tail sheath knockout (ΔTS) or baseplate knockout (ΔBP) strains were inoculated into 50 ml of YEME media at a final concentration of 10^7^ SFU /mL and grown at 30 °C. At **a**, 48 hr, **b**, 54 hr, or **c**, 74 hr post-inoculation, 1 mL samples were collected, washed in saline, and stained with equal proportions of SYTO 9, a membrane permeant nucleic acid stain, and propidium iodide, which stains only dead or damaged cells. Therefore, dead and live cells are visualized by red and green fluorescence, respectively. Samples were imaged using a fluorescent microscope within 30 minutes of staining. The ratio of live/dead cells was calculated from total fluorescence intensity values per image. One-way ANOVA and Dunnett’s multiple comparisons test were performed to calculate statistical significance. Adjusted P-values are shown above the relevant graphs. Diameters of individual mycelial pellets were manually measured from each collected image (images measured for WT, ΔTS and ΔBP, respectively; at 48 hr – n=99, 110, 130, at 54 hr – n= 173, 114, 138 and at 74 hr – n= 73, 70, 71).One-way ANOVA and Dunnett’s multiple comparisons test were performed to calculate statistical significance. Adjusted P-values are shown above the relevant graphs. Representative images are shown here. Experiments were carried out at least three times for each strain.

To address the role of the AAA+ ATPase protein in eCIS function we performed the same experiment as described above using the ΔATPase *Sco* mutant strain. Although this strain produces eCIS particles with a normal appearance, the mycelial pellets produced by this strain after 54 hr of growth in liquid media resembled those produced by the eCIS-deficient mutant strains. These pellets were approximately half the size of the WT pellets, and they displayed half the frequency of cell death (Supplementary Fig. 4b). This trend continued at the 74 hr time point (Supplementary Fig. 4c). As with the other eCIS-deficient mutants, the mycelial pellets of the ΔATPase strain resembled WT after 48 hr of growth (Supplementary Fig. 4a). These data demonstrate that the presence of tail-like particles alone does not mediate cell death and that the ATPase performs a function that is required for the biological activity of the eCIS particles.

### *Sco* strains lacking eCIS display altered development

*Sco* follows a developmental program characterized by a shift from formation of vegetative hyphae to aerial hyphae, which is followed by spore formation. To determine whether the lack of eCIS might perturb this program, we compared the growth of the WT and ΔTS strains on solid media. At early stages of growth, visual examination of the agar plates did not reveal any significant differences, with all strains producing similar vegetative lawns. However, after 48 hours, eCIS mutants had visibly developed faster in terms of producing the pigmented secondary metabolite, actinorhodin (Figure 4a, top panel). At later time points, any visible apparent differences disappeared, and by 72 hours all strains had developed comparable lawns of aerial mycelia (Figure 4a, bottom panel). In liquid media, we observed that WT and eCIS-deficient strains exhibited similar growth rates (Supplementary Fig. 5).

**Fig. 4.**
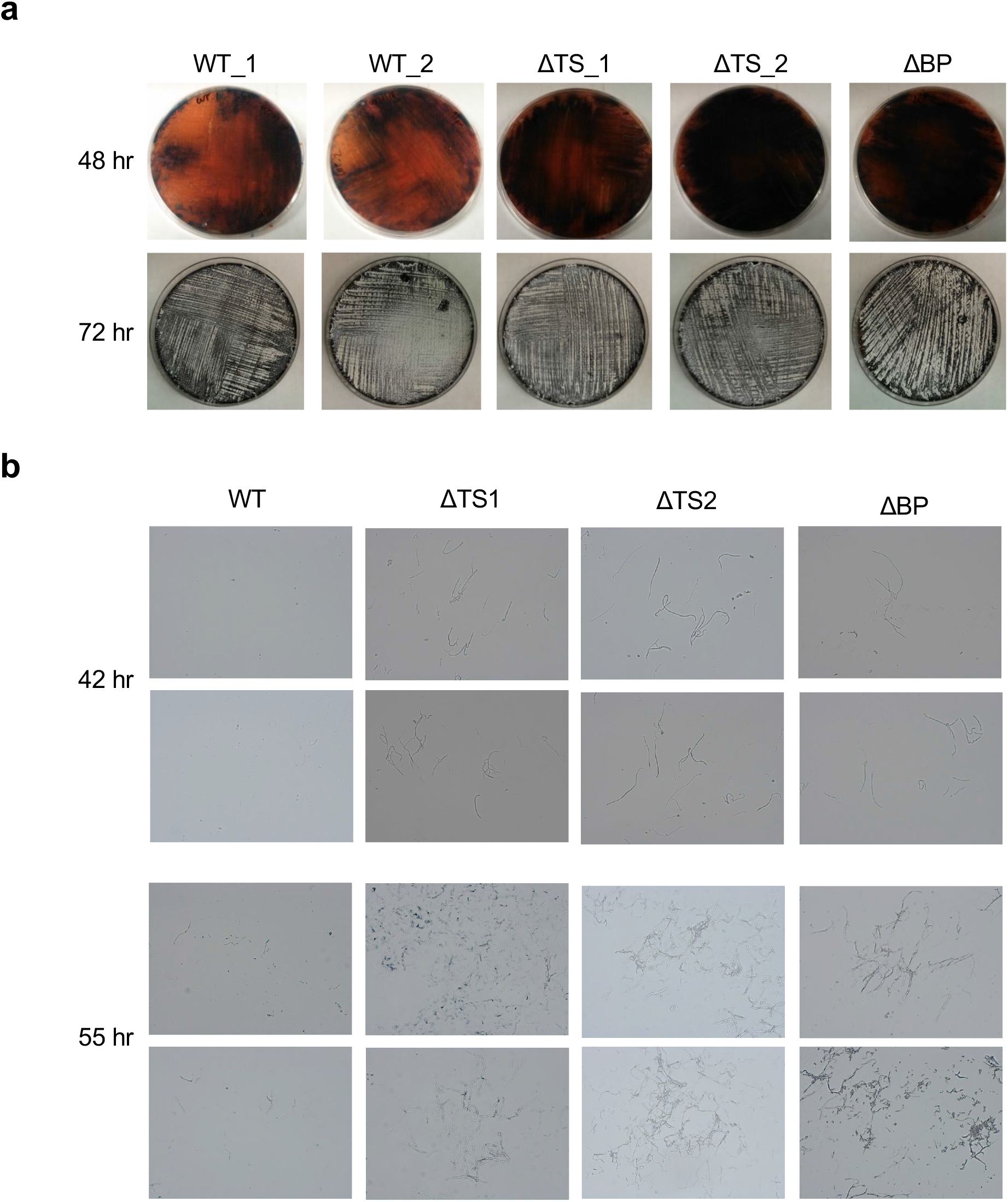
eCIS mutants display accelerated development and early spore formation on solid medium. **a**, 10^8^ SFU of *Sco* wild-type (WT), two independently isolated tail sheath knockout strains (ΔTS_1 and ΔTS_2) or a baseplate knockout strain (ΔBP) were plated on R2YE agar. Top panel shows representative images of secondary metabolite development in lawns grown for 48 hours; bottom panel shows aerial hyphae development on the same plates as above, at 72 hours of growth. **b**, Representative transmission light micrographs of surface imprints on cover slips of agar-grown WT *Sco*, or ΔTS or ΔBP mutant strain lawns 42 and 55 hr post germination. The aerial hyphae and spores stick to the cover slip, but not the vegetative hyphae. Two fields are shown for each sample. Results are representative of three independent experiments with two biological replicates per strain.

It should be noted that the production of actinorhodin by eCIS-deficient strains was highly variable depending on the media used. In some conditions of liquid or solid media growth, these mutants produced considerably less pigment than WT (Supplementary Fig. 6a,b,c). Since the regulation of actinorhodin production is very complicated^26–30^, it is difficult to rationalize these variable effects. However, the perturbation of secondary metabolite production in the absence of eCIS is consistent with a role for it in the developmental program.

We used a microscopy approach to gain a clearer assessment of aerial hyphae formation at intermediate time points in the developmental program. We applied glass cover slips to the surface of the growing bacterial lawn. Since the surface of the cover slips is hydrophobic, only aerial hyphae, which have a hydrophobic coating, stick to them. Adhered hyphae were then detected through microscopic examination. No cells were seen on cover slips that had been applied to the surface of lawns in any of the samples up to 35 hr post-germination. At 42 hr, the WT strains had yet to produce any visible aerial hyphae, while the ΔTS and ΔBP mutant strains showed well-developed aerial hyphal strands clustered in numerous regions on the microscope slide (Fig. 4b). This trend continued at 55 hr of growth where the WT strains had developed a few dispersed hyphal strands compared to the eCIS-deficient mutants, which displayed larger, more developed clusters of hyphae (Fig. 4b).

### eCIS-encoding genes are developmentally regulated

Since the absence of eCIS appeared to perturb the *Sco* developmental program, we wondered if the eCIS-encoding genes were expressed in a developmentally regulated pattern. To this end, we fused four known eCIS region promoters (Fig. 5a; Supplementary Fig. 7), to plasmid-borne genes encoding luciferase activity (*luxCDABE*)^31^. Introduction of these plasmids into WT *Sco* revealed that all four eCIS promoters were active and displayed the same temporal pattern of expression. This pattern was similar to the promoter of the major sigma factor gene *hrdB*^32^, which acts during vegetative growth and to the promoter of *redD*. This gene encodes the transcriptional regulator of the *red* biosynthetic cluster, which is transcribed at early stages of the metabolic switch^33,34^. These results show that the eCIS operons are expressed during the vegetative growth phase under normal growth conditions, right before the metabolic switch that leads to aerial hyphae formation and sporulation.

**Fig. 5.**
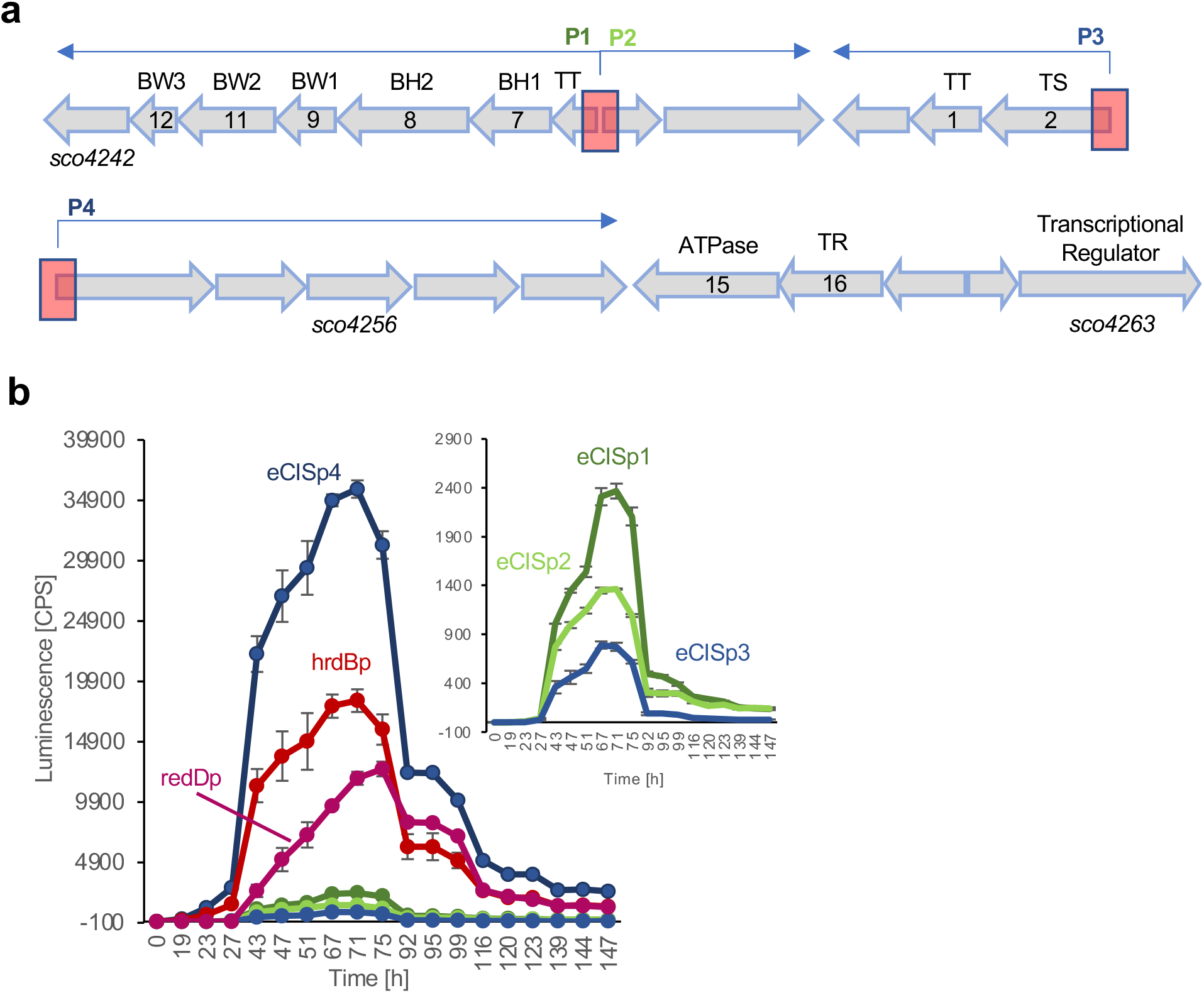
The eCIS cluster of *S. coelicolor* is expressed during vegetative growth right before the metabolic switch. **a**, Schematic representation of the eCIS cluster of *Sco*. Open reading frames and their transcriptional direction are represented by block arrows and are drawn to scale. Gene name is shown under the first and last genes in the cluster, spanning from *sco4242-sco4263*. The encoded eCIS function is shown as an abbreviation above each ORF. The Afp eCIS annotation, which is used in some other publications, is shown inside the arrows (Afp1-16^18^). The two divergent promoter regions are shown as red blocks. Forward and reverse transcriptional direction shown in blue arrows above the cluster. **b**, A fusion construct of each eCIS promoter (eCISp1-p4, corresponding to promoters P1-P4 shown in (a)) with *luxCDABE* was conjugated with *Sco* and grown on 96-well MS agar plates for 7 days. *hrdBp* and *redDp* reporter strains were used as controls. Luminescence was measured three times a day. Activity of the weaker promoters: eCISp1, eCISp2 and eCISp3 is shown in enlarged view in the top right insert. Three biological replicates were tested for each assay (n = 3). The mean normalized to empty vector is shown with error bars representing standard deviation of the mean.

Our observed transcriptional pattern for the eCIS promoters is consistent with previous observations that eCIS expression is regulated by *bldA*^35–37^. This gene is required for proper morphological development, secondary metabolism, and sporulation^38–40^. It encodes the only tRNA for the rare leucine codon, TTA, present in 2-3% of *Streptomyces* genes. The production of the eCIS transcriptional regulator encoded by *sco4263* is dependent on BldA as it contains a TTA codon. We analyzed 56 *Streptomyces* eCIS regions and found that 42 contained at least one gene containing an in-frame TTA codon. Of the identified TTA-containing genes, 22 are putative eCIS regulatory genes (Supplementary Data Sheets 1 and 2). These findings indicate that eCIS regions are often regulated by *bldA*, which ensures that eCIS particles will accumulate in conjunction with the morphogenetic transition.

### Identification of a putative toxin and fiber for *Streptomyces* eCIS

To address the question of whether eCIS may perform a shared function among many *Streptomyces* species, we sought to identify conserved features particular to *Streptomyces* eCIS. To this end, we searched publicly available *Streptomyces* genomes and identified 153 eCIS-encoding regions within 127 *Streptomyces* genomes (Supplementary Data Sheet 3). eCIS have been classified previously into 6 distinct subtypes based on sequence similarity and gene arrangement^3^. The majority (> 80%) of *Streptomyces* species including *Sco* encode a type IId eCIS, as has been previously noted^3^; thus, we assume that any conserved eCIS function must be performed by this type.

Most eCIS regions encode proteins with domains associated with toxin activity. These proteins are thought to be assembled into the eCIS particle and injected into target cells to mediate cell killing^10,41,42^. We were not able to identify a protein encoded frequently within *Streptomyces* eCIS regions that contained an identifiable toxin domain. However, a BLAST^43^ search with the Sco4256 protein, which we found to be a component of the *Sco* eCIS particle (Supplementary Table 1), revealed closely related proteins encoded in 90% of the 105 *Streptomyces* class IId eCIS (Supplementary Data Sheet 4). Alignment of a representative group of sequences and structural analysis using HHpred^44^suggested a three domain architecture with the second domain predicted to be a transmembrane helical region (Fig. 6a, Supplementary Data Sheet 5). The first domain, present across all homologues, shows high probability similarities to periplasmic chaperones involved in pilus assembly among other structures and families (Supplementary Data Sheet 5). Approximately one third of homologues, including the one from *Sco*, possess a C-terminal third domain with similarity to various bacterial toxins and hemolytic lectins (Fig. 6b; Supplementary Data Sheet 5). These proteins may be delivered by eCIS as toxic cargo. Interestingly, homologues of these proteins were also identified in class IId eCIS regions in genomes of other Actinobacterial species and several species of filamentous Cyanobacteria and Chloroflexi (Fig. 6a; Supplementary Data Sheet 4). Proteins with highly significant similarity to Sco4256 were rarely found in non-eCIS encoding regions.

**Fig. 6.**
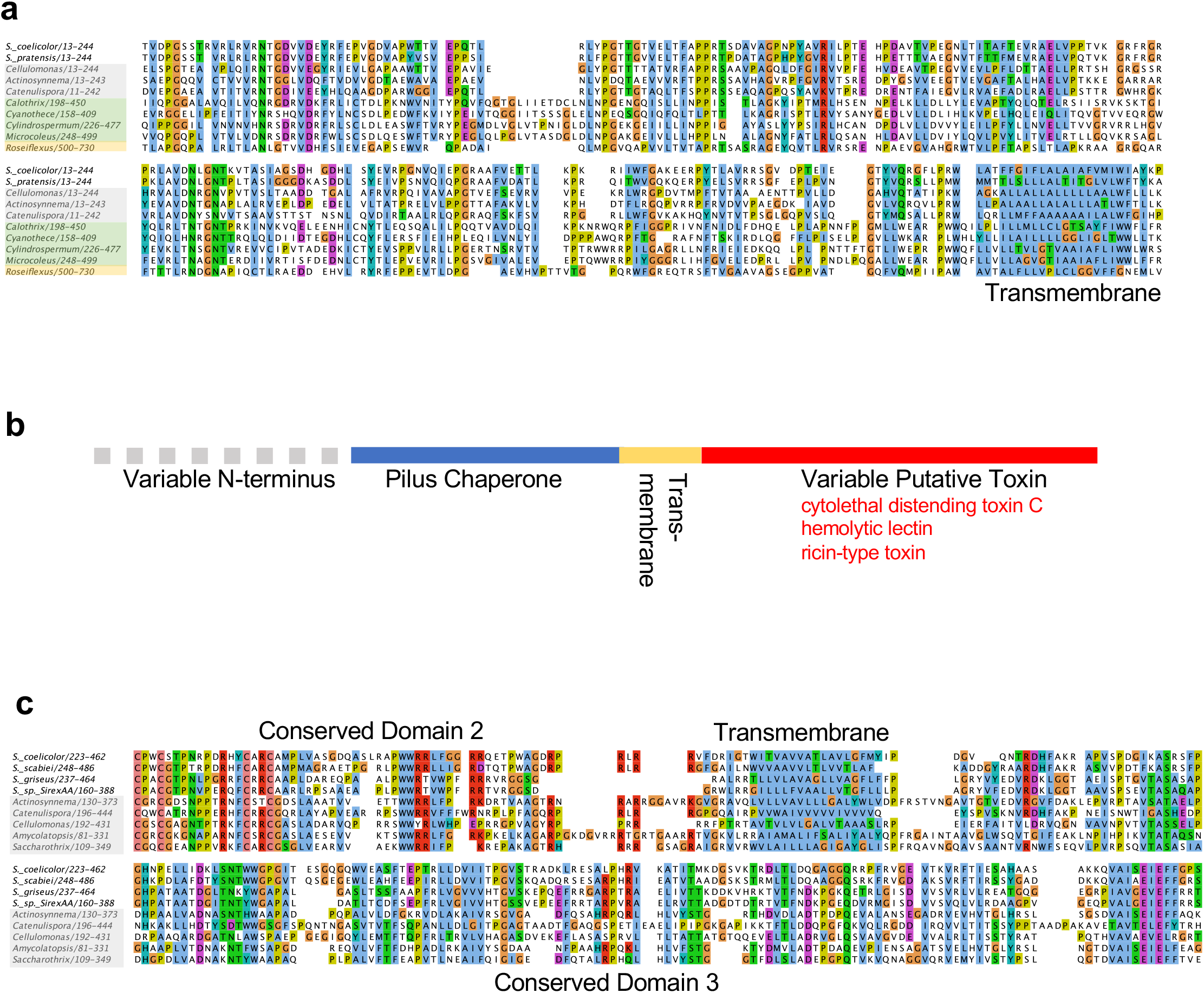
Conservation of the putative toxin and fiber of the *Sco* eCIS. **a**, A diverse selection of hits from a BLAST search initiated with Sco4256, the putative toxin, were aligned and analyzed by HHpred. Sequences shown are from *Streptomyces* species (unshaded), other Actinomycete species (shaded gray), Cyanobacteria (shaded green), and Chorflexi (shaded yellow). All proteins are associated with type IId systems and all genera are filamentous except for *Cyanothece*. **b**, A schematic of the conserved domains of proteins related to Sco4256 is shown. **c**, A diverse selection of hits from a BLAST search initiated with Sco4242, the putative fiber, were aligned. Sequences shown are from *Streptomyces* species (unshaded), other Actinomycete species (shaded gray). These proteins are found in type IId eCIS. All alignments were performed with MUSCLE^59^ as implemented in Jalview^60^. The clustal coloring scheme was used.

Structurally characterized eCIS^5,13,19^ and contractile phage tails^1^ have fiber proteins attached to their baseplates. In phages, these fibers mediate host cell attachment. In most contractile-tailed phage genomes and in the genomes of characterized eCIS^8,11,13,14^, fiber proteins are encoded immediately downstream of genes encoding the baseplate wedge proteins, BW2 and BW3. In the *Sco* eCIS region, the protein product of the gene in this position (*sco4242*) (Fig. 5a) was found to be a component of the eCIS particle (Supplementary Table 1). A BLAST search with this protein revealed closely related homologues in 75% of class IId eCIS regions in *Streptomyces*, and in the class IId eCIS of other Actinomycete species (Fig. 6c; Supplementary Data Sheet 6). Alignment of these proteins revealed a conserved C-terminal region approximately 150 residues long, which is preceded by a strongly predicted transmembrane motif (Fig. 6c). The C-terminal domain has strong similarity to carbohydrate-degrading domains, as predicted by HHpred (Supplementary Data Sheet 7). Since receptor binding proteins of phages, of which fibers are one group, often bind to or degrade carbohydrates on the bacterial cell surface, these predicted functions of the Sco4242 protein are consistent with its serving as a bacterial cell surface binding protein. Conserved domain 2, comprising approximately 50 residues immediately preceding the transmembrane segment is Cys-rich and may be a Zn-binding domain as predicted by HHpred (Supplementary Data Sheet 7). This region might mediate attachment of these proteins to the baseplate. In summary, our bioinformatic data (summarized in Supplementary Data Sheet 1) show that most *Streptomyces* type IId eCIS share related putative fiber and toxin proteins, supporting a common function for eCIS in this genus.

## Discussion

eCIS particles are produced by bacterial species in widely divergent bacterial clades. The dispersed occurrence of eCIS and their demonstrated roles in mediating toxic activities against eukaryotic cells has led to a view of these systems as defense mechanisms, useful under specific conditions but not central to normal cellular function. The results presented here introduce an entirely distinct role for eCIS in *Sco* where they appear to participate in the developmental process of this species. In particular, we have shown that *Sco* produces eCIS particles as part of its normal growth cycle and that strains lacking functional eCIS particles exhibit pronounced alterations in their developmental program. These changes included perturbation of antibiotic production during growth on solid and in liquid media (Fig. 4a; Supplementary Fig. 6), and accelerated development of aerial hyphae on solid media (Fig. 4b). Most intriguingly, eCIS-deficient mutants displayed significantly reduced levels of cell death and altered morphology during liquid growth (Fig. 3). These results suggest that *Sco* eCIS function by inducing intra-strain lethality, which may play a role in the developmental process.

In contrast to a previous study on *Streptomyces lividans*, we did not detect eCIS mediated inhibition of *S. cerevisiae* growth by *Sco*^21^. Given that the eCIS systems in these two species are very closely related, we would expect them to perform the same function. It is possible that we did not detect the growth inhibitory activity of the *Sco* eCIS because we used a different assay that may not be as sensitive as that used in the other study. In addition, since we observed complicated changes in antibiotic production by the eCIS-deficient strains, it is possible that indirect effects on antibiotic production caused the effects seen in the previous study. This idea is supported by the absence of *S. cerevisiae* growth inhibition induced by strain M1152 (Fig. 2), which does not produce the four major *Sco* antibiotics.

While intracellular Sco lysates displayed mostly fully sheathed eCIS structures with typical appearance (Fig. 1a), extracellular samples almost exclusively contained empty contracted sheath species (Fig. 1c). This stark difference between the intracellular and extracellular pools of eCIS-derived structures suggested that eCIS were not escaping from cells through generalized lysis, a conclusion also supported by the absence of ActR, a cytoplasmic protein, in the extracellular fractions containing sheath proteins (Supplementary Fig. 1b). A model that could account for the above observation is that in intact mycelia the tail-like structures are membrane-bound. Upon encountering the appropriate stimulus, the eCIS may extend through the membrane to contact a neighboring cell, resulting in contraction and release of the contracted sheath. The possibility of the *Sco* eCIS being attached to the cell membrane is supported by the conserved presence of a strongly predicted transmembrane helix in its putative fiber protein (Fig. 6c). In recent structural studies on the eCIS of *Anaboena*, a filamentous cyanobacterium, the eCIS structures were shown to be anchored to the thylakoid membrane by a transmembrane helix in a fiber-like protein that was attached to the baseplate^13^. Although we did not observe fibers on the *Sco* eCIS particles, we did detect the putative fiber protein in our purified samples by mass spectrometry (Supplementary Table 1). The conditions of cell lysis during purification would likely disrupt membrane interactions leaving the fibers in variable conformations that might be difficult to distinguish by TEM.

One potential limitation of our study is that some key experiments, such as those investigating cell death, were performed in liquid cultures in which *Sco* does not sporulate. However, the developmental program with respect to gene expression has been shown to be similar in liquid and solid medium^45,46^. Thus, we expect to see related differences during growth on solid media where cell death has also been shown to occurs^47–49^.

Vegetative hyphal degradation during development has long been recognized as a feature of *Streptomycete* development^50^ and more recently programmed cell death has been postulated to occur as part of the shift from vegetative to aerial mycelial growth^24,47,51^. It is thought that a coordinated stage of cell death limited to a subset of the vegetative hyphal population during development may serve to provide additional nutrients to the growing reproductive aerial mycelium and dead hyphae may also serve a structural role or facilitate nutrient transport ^48,49,52,53^. Consistent with a role at this stage in the *Sco* lifecycle, we have shown that eCIS expression peaks late in the vegetative growth phase in a temporal pattern similar to the antibiotic regulator gene, *redD*. The requirement of BldA, a master regulator of the morphogenetic switch, for the expression of the *Sco* eCIS genes also supports a role in this process. The apparent requirement for *bldA* for expression of eCIS genes in many *Streptomyces* species as well as the conservation of putative effector and fiber proteins among many *Streptomyces* eCIS regions suggest that these eCIS may share a conserved function in programmed cell death. This shared function in the *Streptomyces* lifecycle would explain the very frequent occurrence of eCIS in this clade.

Interestingly, a role in intra-strain cell death was recently proposed for the eCIS of *Anabaena*, a filamentous cyanobacteria with a complex lifestyle^13^. We found that the eCIS of diverse Actinomycetes and other filamentous Cyanobacteria encode homologues of the Sco4256 putative effector protein, providing another connection between eCIS function in these two filamentous groups of bacteria.

To conclude, this study has revealed a previously unknown function for eCIS in mediating intra-strain cell lethality that impacts the developmental process of *Streptomyces*. The possibility that this function is conserved among diverse species opens up exciting avenues for future research.

## Methods

### Bacterial strains, plasmids, and media

Bacterial strains, plasmids and cosmids used in this chapter are listed in Supplementary Table 3. All *Streptomyces* strains used were derivatives of *Streptomyces coelicolor* A3(2). *Sco* strains were grown at 30 °C on maltose-yeast extract-malt extract (MYM) agar medium^54^ for 3 days or in yeast extract-malt extract (YEME) liquid medium^55^ with continuous agitation at 180 rpm for 3 days, unless indicated otherwise. Other bacterial strains were grown in Lysogeny Broth (LB) medium or agar (pH 7.0) at 37 °C unless indicated otherwise. *Saccharomyces cerevisiae* were grown in YPD medium or agar (pH 7.0) at 30 °C.

### Construction and testing of Lux reporter strains

The eCIS promoter sequences were amplified from wild-type *Sco* M145 genomic DNA using primers SCO_eCISp1p2GEN and SCO_eCISp3p4GEN (Supplementary Table 4). The resulting PCR fragments were used as a template to introduce an EcoRV blunt cut site using SCO_eCISp1p2EcoRV and SCO_eCISp3p4EcoRV primers. The PCR products were cloned into the EcoRV site of the pF*lux* plasmid^31^ both in the forward and the reverse orientation for the ability to test both transcriptional directions of the divergent promoter regions. All plasmids were verified by sequencing. The resulting reporter plasmids pF*lux*-eCISp1-p4 were electroporated into methylation-deficient *E. coli* ET12567/pUZ8002 cells^55^. Plasmids were introduced into wild-type *S. coelicolor* by intergenic conjugation and selection on apramycin (50 *μ*g/mL)^55^.

For luminescence assays 10^4^ spore forming units (SFU) of each *Sco* reporter strain were resuspended in 10 *μ*L of saline and plated in triplicate onto 200 μL of soya-mannitol (MS) agar^56^ in 96-well polystyrene plates. Plates were incubated for 7 days at 30°C and luminescence was measured three times daily using a PerkinElmer Victor X multilabel plate reader.

### Generation of gene replacement mutants

To create genomic replacements of the eCIS tail sheath, baseplate and ATPase encoding genes the REDIRECT method for PCR targeting in *Streptomyces* was employed as described^55^. The *aac3(IV)-oriT* cassette which confers resistance to apramycin, was amplified using SCO4253_Disrp, SCO_BP_Disrp or SCO4259_Disrp primer pairs, respectively (Supplementary Table 4). The amplified cassettes were used to replace the respective eCIS genes in cosmid StD8a^57^. The resulting cosmids (listed in Supplementary Table 3) were introduced by conjugation from the non-methylating *E. coli* strain ET12567/pUZ8002^55^ into the wild-type strain M145 to obtain apramycin-resistant exconjugants. To select for double cross-over exconjugants the aparamycin resistant colonies were screened for kanamycin resistance. Three Apra^R^ Kan^S^ exconjugant strains were selected for each gene, resulting from separate conjugation events. Mutant strains were confirmed by PCR using oligonucleotides; StD8aΔTS_aac(3)IV, StD8aΔBP_aac(3)IV or StD8aΔATP_aac(3)IV (Supplementary Table 4), respectively and by DNA sequencing.

### Actinorhodin production assay

Wild-type *Sco* M145 or *Sco* Δ*sco4253* (ΔTS) were inoculated in experimental triplicate into R2YE liquid media at a final concentration of 1.5×10^6^ SFU/mL. Cultures were grown with agitation for 22 hours at 30 °C. All cultures were standardized to an OD_450_ of 0.5. Cells were recovered by centrifugation at 3500 x g for 10 min, resuspended in saline and centrifuged again at 3500 x g for 10 minutes. Cell pellets were resuspended in 50 mL of fresh RG2 minimal media. The cultures were incubated at 30 °C with continuous shaking for 94 hr. For visual comparison of pigment production, images of the growing cultures were taken using a NIKON D3000 camera at a fixed position within a light box at times of 0, 18 hr, 24 hr, 41 hr, 49 hr, 65 hr, 70 hr and 94 hr. For quantification of total actinorhodin production, 480 *μ*l samples were collected at the time points indicated above. 120 *μ*l of 5M KOH was added to a final concentration of 1M, and samples were vortexed and centrifuged at 3000 x g for 5 minutes. The absorbance of the supernatant at 640 nm was read using a polystyrene 96 well plate. Each assay was standardized to the weight of the pellet.

### Aerial mycelium formation assay

10^8^ SFU of *S. coelicolor* wild-type or eCIS-deficient mutant strains in biological duplicate were plated on R2YE agar plates and grown at 30 °C. Starting from 24 hr post germination, sterile glass cover slips were gently applied to the top surface of each bacterial lawn at approximately 12-hour intervals. Cover slips were sealed on top of a microscope glass slide and examined using a Zeiss transmitted light microscope at 40 X magnification.

### Inter-species growth inhibition assays

Spores (10^5^ SFU suspended in 5 *μ*L of 0.85% saline) of wild-type *S. coelicolor* M145, M1152 or the eCIS mutant strains ΔTS or M1152 ΔTS were spotted on R2YE agar plates. Plates were incubated for 3 days at 30°C before being overlaid with 5 mL molten, extra-soft LB agar (0.5% agar) containing organisms to be tested (at a final OD_600_ ranging from 0.01-0.1). Plates were incubated at 37 °C overnight and antimicrobial activity was assessed by measurement of zones of clearance in the indicator lawn. Assays against *Streptomyces* strains were conducted by overlaying a 3-day colony plate with 1 mL 0.85% saline solution containing 5 *μ*L of concentrated *Streptomyces* spores. Plates were incubated for 3 days at 30 °C before antimicrobial activity was assessed. Assays against *Saccharomyces cerevisiae* were conducted by overlaying a 4-day colony plate with 5 mL molten, extra-soft YPD agar containing *S. cerevisiae* (at a final OD_600_ ranging from 0.01-0.05). Plates were incubated at 30 °C overnight.

### Generation of antibodies against TS and BW1 proteins

The TS and BW1 proteins were expressed and purified from *E. coli*. Genes encoding the BW1 protein (*sco4245*) and the TS protein (*sco4253*) were cloned from *Sco* genomic DNA into the BseR1 site of p15TV-L vector using the In-fusion HD cloning kit (Clonetech) using primers listed in Supplementary Table 4. Verified plasmids were transformed into *E. coli* strain BL21 (DE3), grown at 37 °C with appropriate antibiotics to an OD_600_ of 0.6 and protein expression was induced with 1 mM IPTG. Cultures were grown for 16-18 hours at 17 °C. Cells were collected by centrifugation and resuspended in binding buffer (20 mM Tris-HCl pH 7.5; 200 mM NaCl; 5 mM imidazole, 5 mM β-mercaptoethanol (βME)). Cells were lysed by sonication. Lysates were cleared by centrifugation at 17,000 rpm for 20 minutes at 4 °C. Proteins were affinity-purified by incubation with 2 mL of Ni-NTA agarose resin (Invitrogen) for 30 minutes at 4 °C. The mixture was passed through a column at room temperature and washed extensively with binding buffer containing 30 mM imidazole. Bound proteins were eluted with binding buffer containing 300 mM imidazole and dialyzed overnight at 4 °C in buffer containing 20 mM Tris·HCl, pH 7.5, 200 mM NaCl, 5% (vol/vol) glycerol, 1 mM DTT. Samples were concentrated using Vivaspin filter concentrators (Sigma-Aldrich) and further separated using Superdex200 16/60 size exclusion column on an ÄKTA fast performance liquid-chromatography system (GE Healthcare). Relevant fractions were analyzed and verified by SDS-PAGE and concentrated to a final concentration of 0.1 - 0.25 mg/ml.

Mouse polyclonal antibodies were generously generated by the Gray-Owen lab at the University of Toronto. Fresh purified protein at a concentration of 0.05-0.25 mg/mL was mixed with 100 *μ*l Emulsigen adjuvant and injected into mice. A second booster dose was administered 21 days later. Antisera were collected on day 35. Antisera specificity was tested by Western blotting with appropriate controls.

### eCIS particle purification

*Sco* strains were grown on MYM agar at 30 °C for 4 days. 1 cm^2^ squares of agar containing mycelium and spores were excised from the plate and directly transferred into 50 mL YEME liquid medium. Cells were incubated at 30 °C for the indicated various timeswith shaking. Extracellular sample (E) was taken from the supernatant. Pellets were washed in 10% sucrose and centrifuged again, resuspended in 16 mL of buffer P^58^ supplemented with 2 mg/mL lysozyme and incubated at 30 °C for 1 hour. Cells were spun down and resuspended in Buffer C (50 mM Tris pH 7.5, 150 mM NaCl, 15% vol/vol glycerol, 1 mM DTT, protease inhibitor cocktail (Sigma-Aldrich-one tablet per 50 mL of buffer)), and sonicated. Lysed cells were centrifuged at 12,000 rpm for 20 minutes and the supernatant was then filtered through a 0.45 *μ*m filter. Filtered samples were spun at 150,000 × g for 3 hours at 4 °C. The pellet was resuspended in 5 mL of ice cold 1XPBS. The samples were loaded onto columns containing 5 mL of DEAE Sepharose anion exchange resin (Sigma-Aldrich), washed extensively with PBS and eluted using 0.5M NaCl in PBS buffer. Samples were ultracentrifuged at 150,000 × g for 90 minutes at 4 °C and pellets were resuspended in 200 μL ice-cold PBS and stored at 4 °C for up to two weeks.

### Western blotting

Samples were mixed in a 1:1 ratio with 2 X SDS loading buffer (100 mM Tris-HCl, pH 6.8; 4% SDS; 20% glycerol; 200 mM β-mercaptoethanol (βME); 0.2% Bromophenol blue). Proteins were separated on a 12% polyacrylamide SDS Tris-Tricine gels and transferred to a nitrocellulose membrane. The membrane was blocked for 1 hour in 5% skim milk in Tris-buffered saline with 1% Tween (TBS-T). When tested with α-eCIS specific antibodies antiserum was added at 1:5000 dilution in TBS-T with 5% milk. Alternatively, α-FLAG primary antibody (Sigma-Aldrich) was added at 1:10,000 dilution. Membranes were incubated overnight at 4 °C. After washing, the membranes were probed with horseradish peroxidase conjugated goat anti-mouse antibody (Santa Cruz Biotechnology 1:2000 dilution in TBS-T) for 1 hour at room temperature and developed using chemiluminescent reagents (GE Healthcare).

### Transmission electron microscopy

Purified eCIS samples were briefly centrifuged to remove impurities and 5 μL of supernatant was applied onto freshly glow discharged carbon-coated copper grids (CF400-CU, 6 nm, 400 mesh, Electron Microscopy Sciences). Samples were allowed to adsorb for 2 minutes before being blotted off by filter paper and grids were washed twice with filtered distilled water. The grids were negatively stained with 15 μL of 2% uranyl acetate for 15 seconds. Grids were imaged with a JEM-1011 (JEOL USA, INC.), digital CDD camera (5 megapixels XR50S, AMT, USA).

### Live/Dead assay

*S. coelicolor* wild-type or eCIS mutant strains were inoculated into 40 mL of YEME culture medium from fresh spores at a density of 10^7^ /mL and incubated at 30 °C with shaking. Samples of 1 mL were centrifuged for 5 min at 12,000 rpm, washed twice and resuspended in 1 mL of distilled water. LIVE/DEAD Bac-Light bacterial viability kit (L7012; Invitrogen) was used to stain cells. The bacterial suspension was mixed with 3 *μ*l of SYTO 9 and propidium iodide (PI) nucleic acid stains, premixed at a 1:1 ratio. The cell suspension and nucleic acid stains were mixed by vortex and left to stand at room temperature for 10 min in the dark. 10 *μ*l of the suspension was deposited on a clean slide and covered with an 18 mm square cover slip. Images were acquired within 30 min of incubation using a Zeiss Axio imager 2, at an excitation of 488 nm and 568 nm and emission of 530 nm (green) or 630 nm (red). Images were analyzed using Zeiss Zen Blue software and statistical analysis was performed on GraphPad Prism 9.

### Mass spectrometry analysis of purified *Sco* eCIS samples

eCIS samples were purified as described above. Extracellular samples were similarly purified from cell-free supernatants following the initial centrifugation step at 7,500 rpm. Purified samples of 30-150 *μ*g total protein were reduced with DTT, alkylated with iodoacetamide, and subjected to tryptic digest. Liquid chromatography tandem-mass spectrometry spectra were collected on a linear ion-trap instrument (ThermoFisher) (SPARC BioCentre, The Hospital for Sick Children, Toronto, Canada). Proteins were identified using Mascot (Matrix Science, London, UK) and analyzed in Scaffold version 3.0 (Proteome Software Inc., Portland, OR, USA). The cut off for protein identification was set at a confidence level of 95%.

## Supplementary Figures

**Supplementary Fig. 1.**
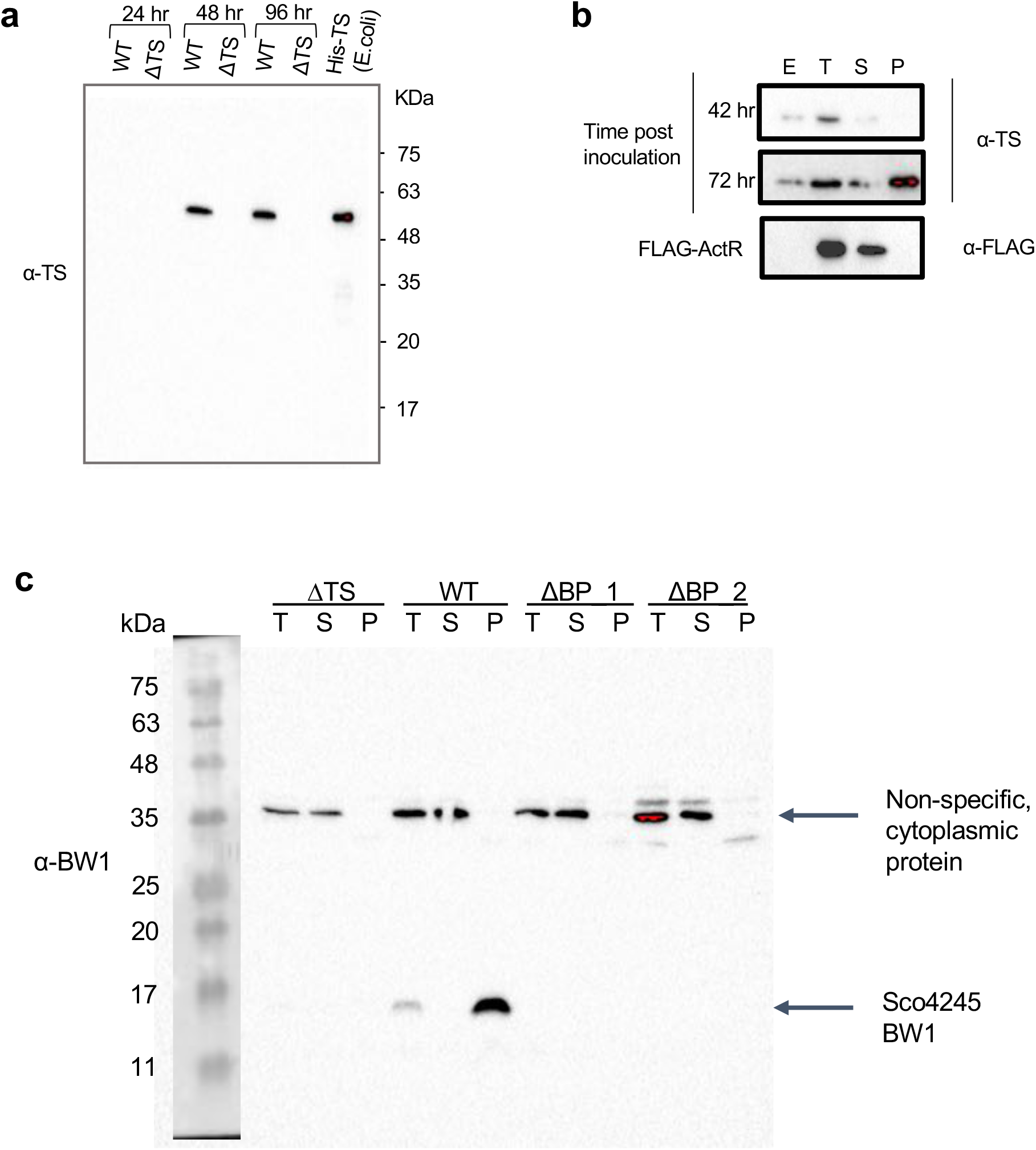
eCIS proteins are detected by specific polyclonal antibodies in lysates of WT *Sco*. **a**, *Sco* wild-type (WT) or a tail sheath knockout strain (ΔTS) were grown in liquid R2YE media to the indicated time points. Whole cell lysates were analyzed by Western blot using α-TS sera. Heterologously expressed and purified 6XHis-tagged-tail sheath (His-TS) was used as a positive control. **b**, WT *Sco* cells were grown in liquid YEME media, cells were collected at the indicated time points and an extracellular “E” sample was taken from the cell-free supernatant. Cells were lysed and an intracellular total lysate “T” sample was collected. The lysate was ultracentrifuged for 3 h at 150,000 X g. Samples of the supernatant “S” and pellet “P” were analyzed. Samples were probed using α-TS antibodies. **c**, *Sco* wild-type (WT) or a baseplate knockout (ΔBP) were grown and fractionated as in (b). Samples were probed by Western blot using α-BW1 antibodies.. For the *Sco* strain expressing FLAG-tagged ActR from a plasmid, expression was induced with 30 *μ*g/mL final concentration of thiostrepton, added 45 minutes before cells were harvested by centrifugation.

**Supplementary Fig. 2.**
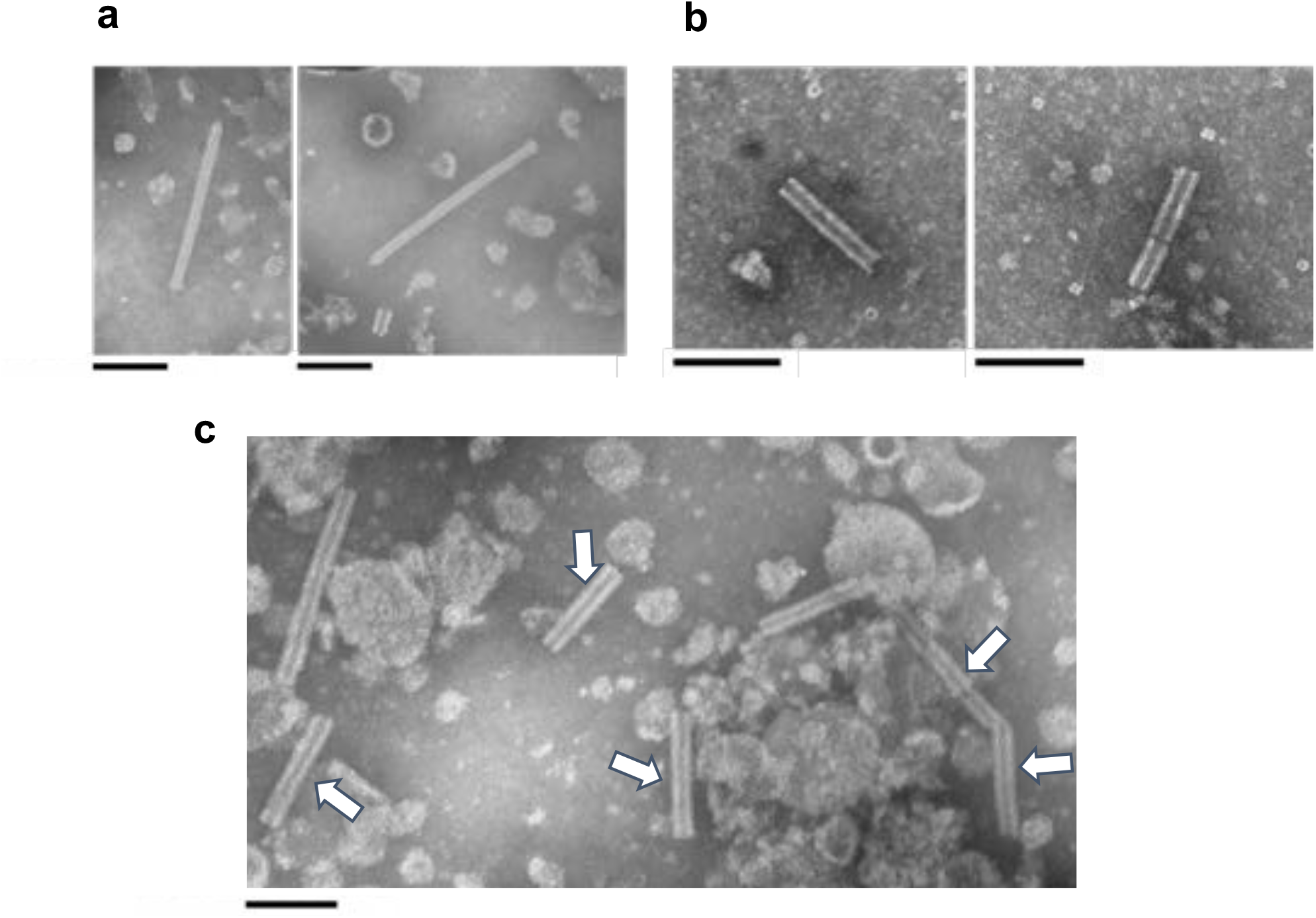
The *Sco* ATPase mutant produces both mature eCIS and empty sheath particles, detectable in the lysate and extracellular media. All images represent *Sco* ΔATPase grown in YEME liquid media. for 48 hr (a), 54 hr (b) or 72 hr (c) post inoculation. **a**, Representative images of purified eCIS particles from cell lysate concentrated to 30 X original culture after 48 hr of growth. The majority of particles at this time are in their extended conformation. **b**, The extracellular fraction after 54 hr of growth is shown. White arrows point to typical emptied contracted tail sheath particles. **c**, Purified lysate from cells grown for 72 hr. Emptied sheath particles are indicated with white arrows. Scale bar represents 100 nm in all images. Images are representative of results obtained for three separate biological replicates.

**Supplementary Fig. 3.**
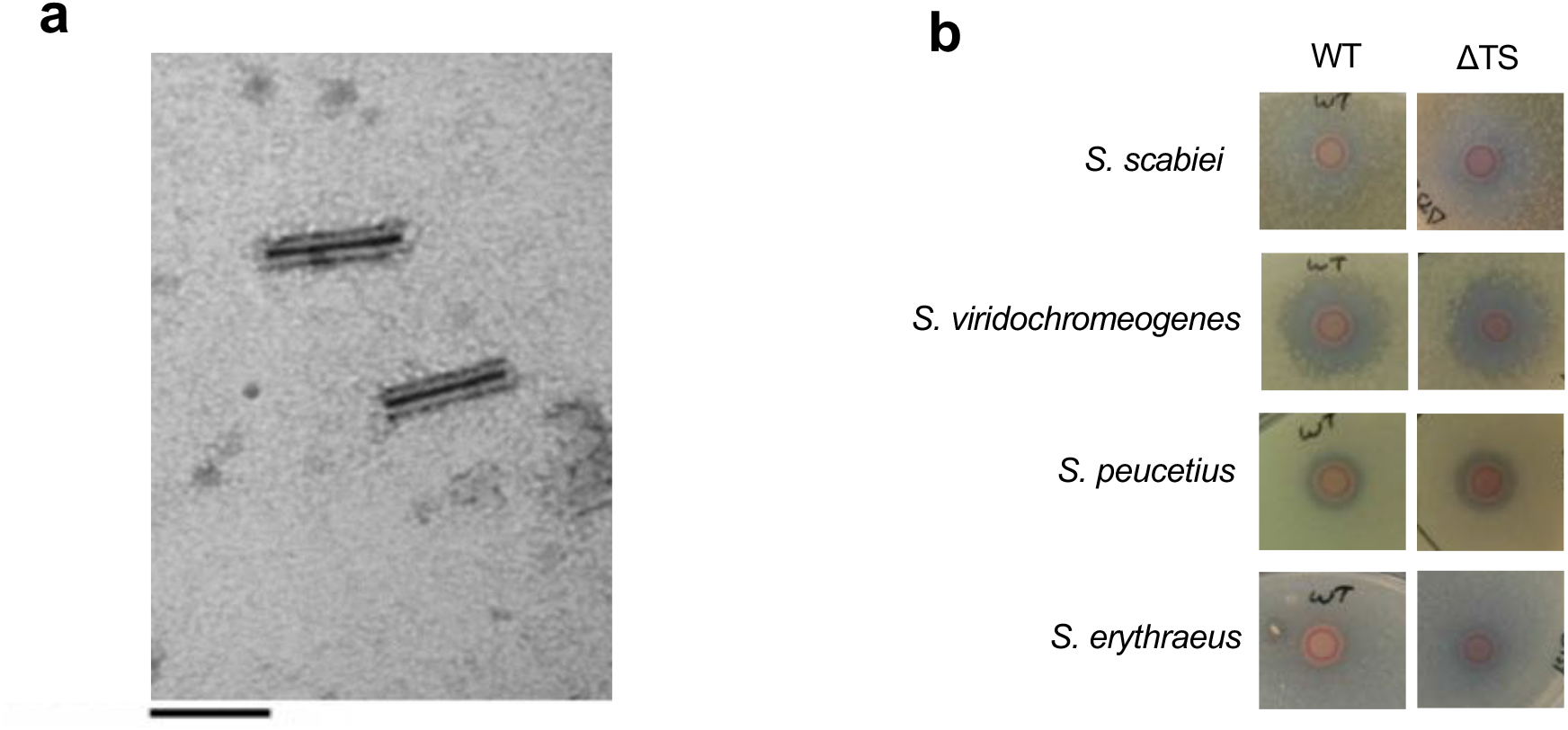
Growth inhibition of other *Streptomyces* species by WT and eCIS-deficient *Sco* strains. **a, A** representative TEM image of extracellular cell-free media collected from *Sco* M1152 strain grown in rich liquid media for 72 hr. Empty contracted sheath particles, visually identical to ones produced by *Sco* WT M145 strain are observed. The presence of these particles is indicative of production of normal eCIS particles that can contract. No particles were detected in a M1152 strain harboring a tail sheath knockout mutation (M1152 ΔTS). Scale bar indicates 100 nm. **b**, Assays were performed by spotting on agar plates 1×10^5^ SFU of *Sco* wild-type (WT) or tail sheath knockout (ΔTS) strains. Cells were grown for 3 days before being overlaid with species to be tested as indicated in each panel. Zones of clearing around the *Sco* colonies indicate lethality or growth inhibitory activity against the indicator lawn. The results shown for a tested species strain were obtained from the same agar plate. Images are representative examples of many replicates (n > 3).

**Supplementary Fig. 4.**
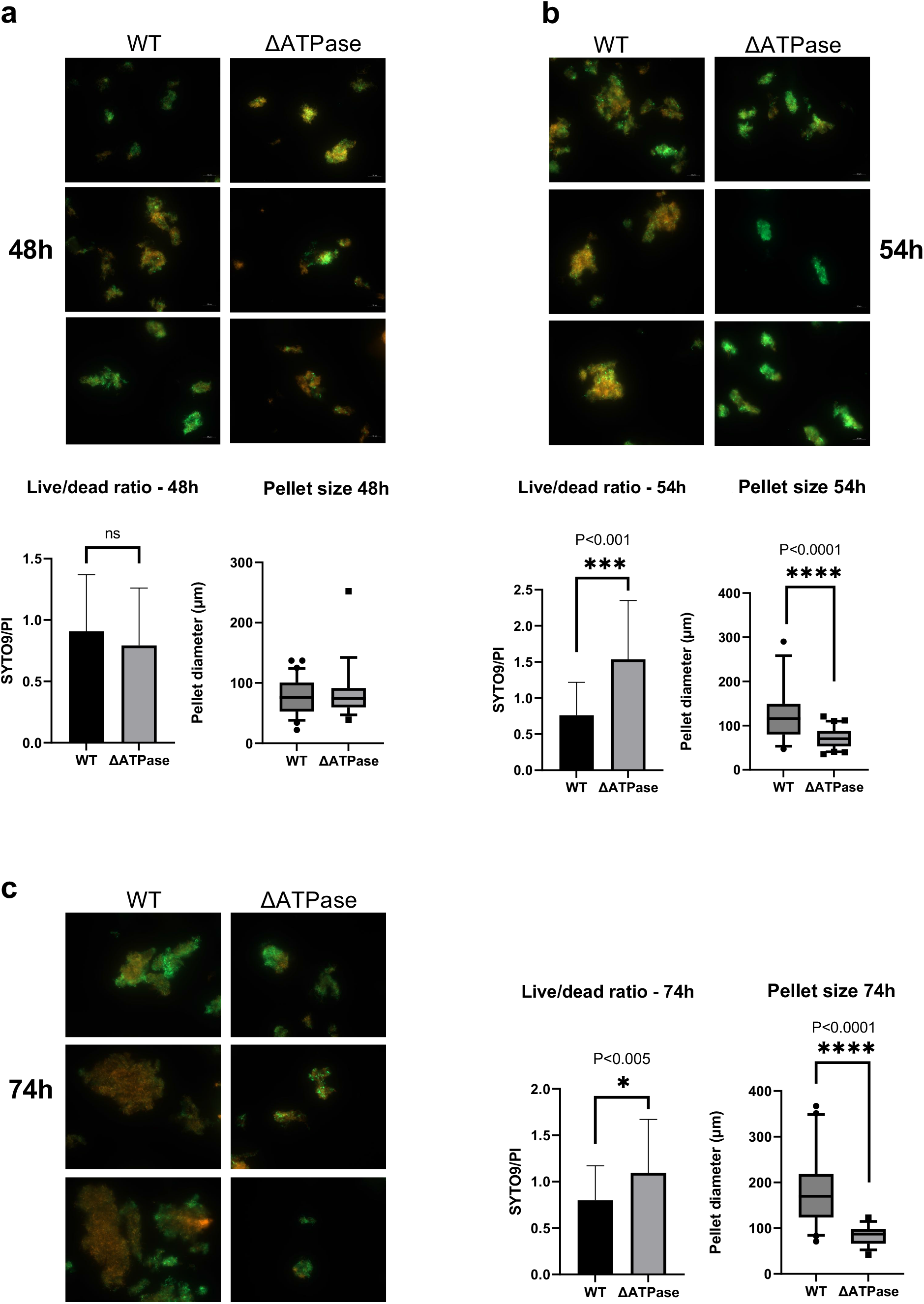
The *Sco* ATPase deletion mutant strain exhibits reduced cell death during hyphal differentiation. **a-c** *Sco* wild-type (WT) or an ATPase knockout (ΔATPase) strains were inoculated into 50 ml of YEME media at a final concentration of 10^7^ SFU /mL and grown at 30 °C. At 48 hr (**a**), 54 hr (**b**) or 74 hr (**c**) post-inoculation 1 mL samples were collected, washed in saline, and stained with equal proportions of propidium iodide and SYTO 9. Total (live and dead) cells are stained with SYTO9 and are visualized by green fluorescence. Cells with damaged membranes (dead cells) are stained only with PI and are visualized by red fluorescence. Samples were imaged within 30 minutes of staining. The ratio of SYTO9 to PI (Live/dead ratio) was calculated from total fluorescence intensity values per image. Diameters of individual hyphal pellets were manually measured from each collected image (n > 30). Single factor ANOVA and Welch’s t-test were performed to calculate statistical significance. Adjusted P-values are shown above the relevant graphs.

**Supplementary Fig. 5.**
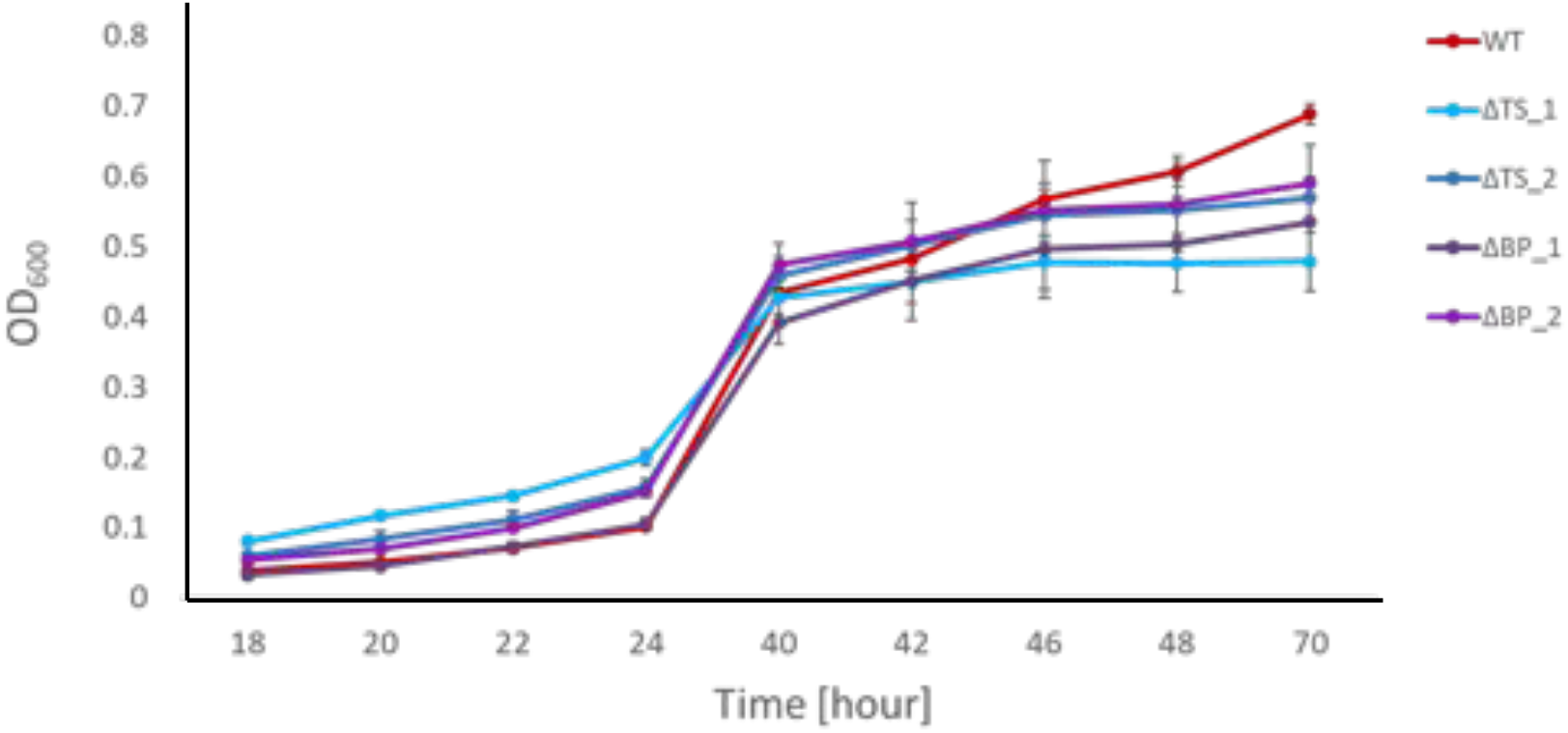
Growth curve for WT *Sco* and eCIS-deficient mutant strains. 10^8^ SFU of *Sco* WT, tail sheath knockout (ΔTS) or baseplate knockout (ΔBP) were inoculated in R2YE liquid media and grown for 70 hours. Optical density (OD_600_) of each sample was taken at the indicated time points. ΔTS_1, ΔTS_2 and ΔBP_1, ΔBP_2 represent two biological replicates of each mutant. The mean is shown with error bars representing standard error of the mean (n=3).

**Supplementary Fig. 6.**
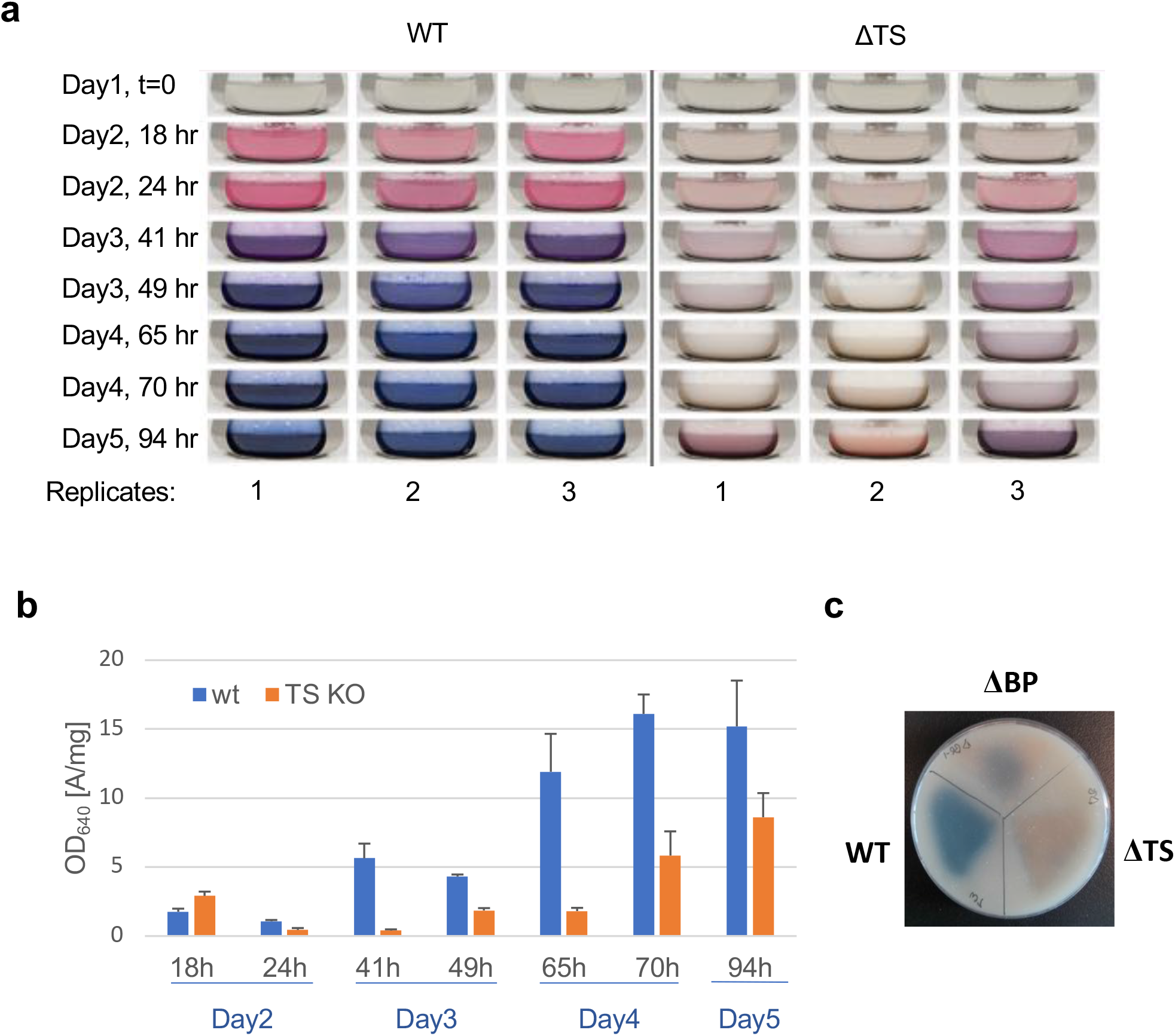
eCIS mutants exhibit decreased actinorhodin production in minimal media. **a**, Comparison of wild-type *Sco* (left) and ΔTS mutant (right) development in liquid media. Three experimental replicates of RG2 liquid media cultures were inoculated to an OD_450_ of 0.5 and incubated with shaking at 30 °C for 94 hours. Images of each culture flask were taken at the indicated time points. **b**, For quantification of total actinorhodin, samples of 0.5 ml were taken at the same time points as in (a). KOH was added to a final concentration of 1 M to extract the actinorhodin and the sample was centrifuged. The optical density (OD_640_) of the supernatant was measured and normalized to wet pellet weight. The mean is shown with error bars representing standard deviation of the mean (n=3). **c**, Equal amounts of *Sco* wild-type (WT), tail sheath knockout (ΔTS) or baseplate knockout (ΔBP) spores were streaked out on minimal Soy Mannitol (MS) agar plates and grown at 30 °C for 4 days. The agar plate shown is representative of multiple independent assays with 3 biological replicates (strains ΔTS 1-3, ΔBP 1-3, each obtained by a separate conjugation event).

**Supplementary Fig. 7.**
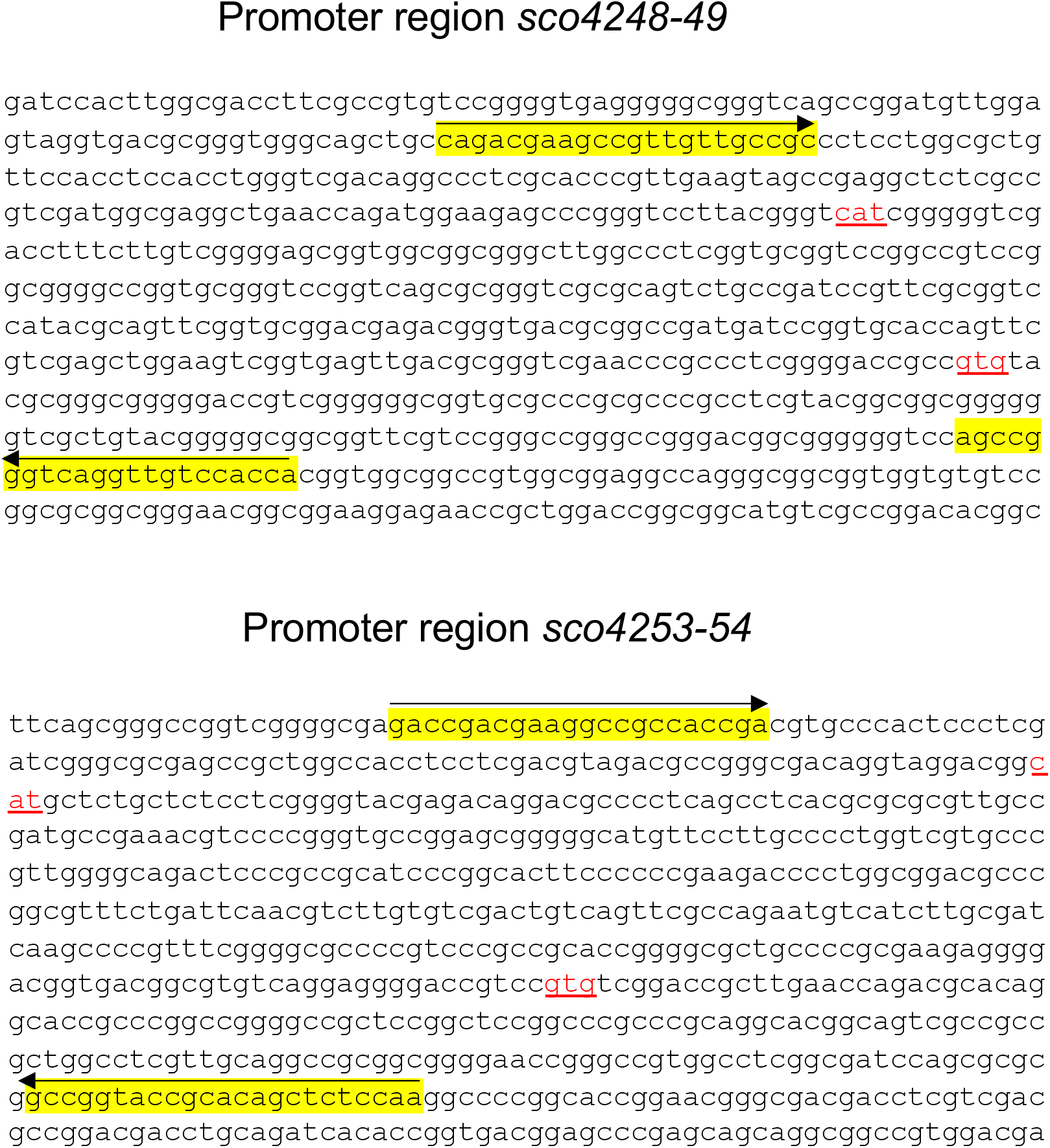
*Sco* eCIS promoter regions. The DNA sequence of the eCIS promoter regions of *Sco* is shown as diagrammed in Fig. 5a. The two divergent promoter regions are indicated by red blocks^37^. The translational start codons (for *sco4248* and *sco4249* on the top and *sco4253* and *sco4254* on the bottom) are underlined in red. Primers used to clone the promoter regions into the *luxCDABE* reporter plasmid are highlighted in yellow and indicated by arrows above the sequence.

## Supplementary Tables

**Supplementary Table 1:**
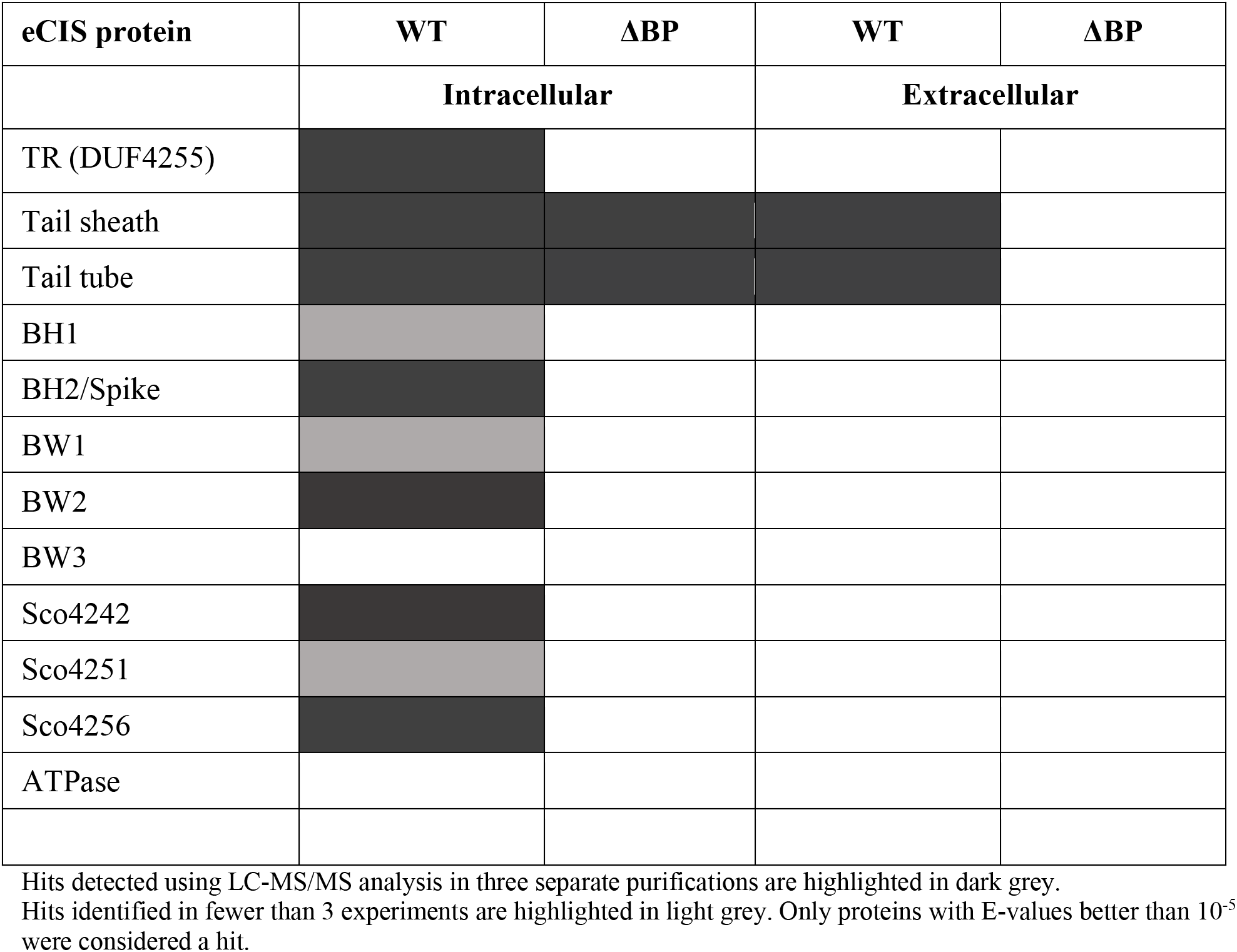
Summary of LC-MS/MS analysis of purified *Sco* eCIS samples.

**Supplementary Table 2:**
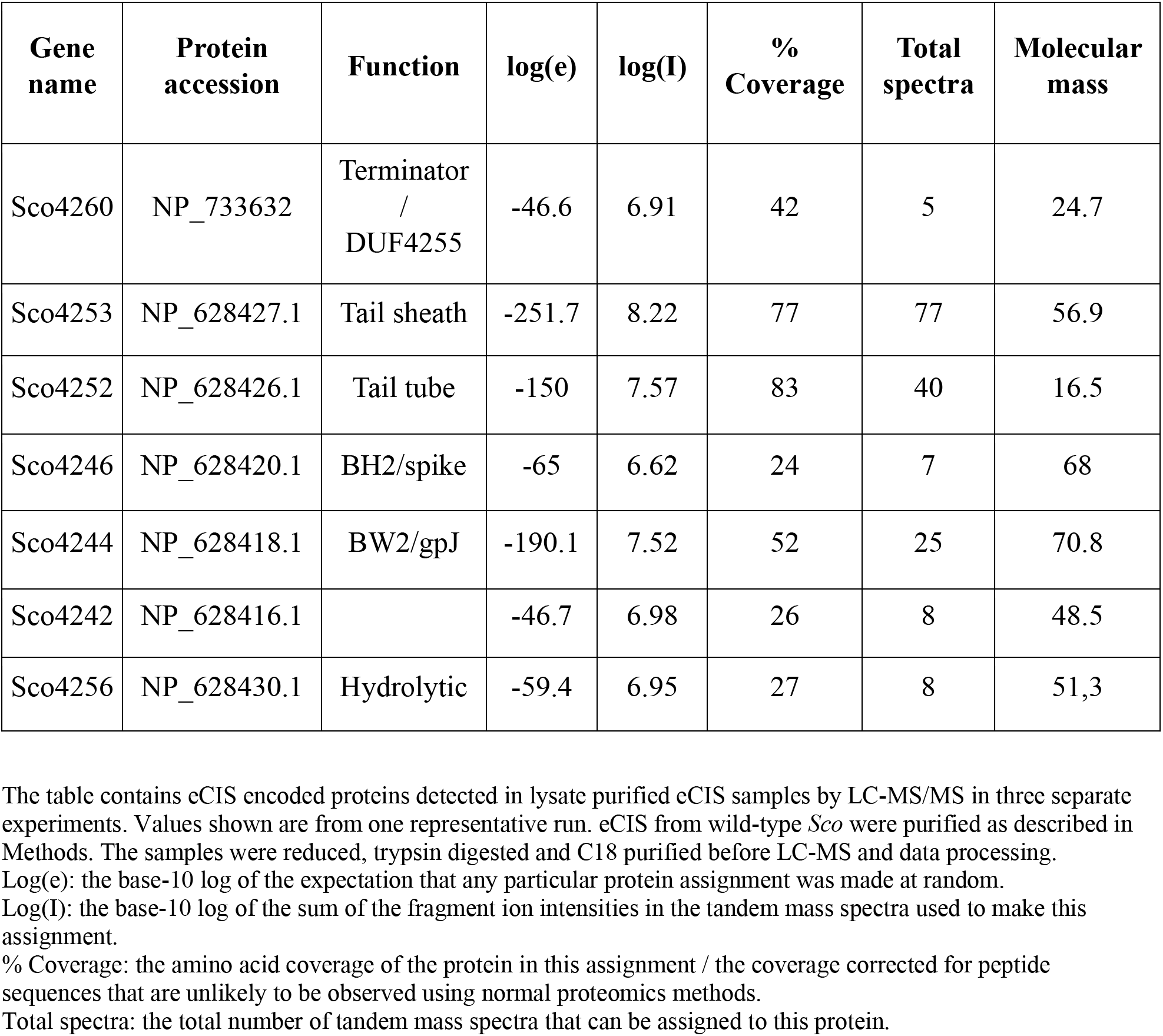
Detailed results of LC-MS/MS analysis of WT eCIS samples.

**Supplementary Table 3:** Strains and plasmids used in this study.

| Strains | Description | Reference or source |
| --- | --- | --- |
| ***Escherichia coli* strains** |  |  |
| DH5α |  | Stratagene |
| ET12567/pUZ8002 | Conjugation donor strain, Kan<sup>R</sup>, Cml<sup>R</sup> | <sup>61</sup> |
| BW25113/pIJ790 | λ Red recombination strain, Cml<sup>R</sup> | <sup>62</sup> |
| BL21(DE3) | Protein expression | Stratagene |
| Stellar HST08 | High transformation efficiency competent cells | Clontech |
| ***Streptomyces coelicolor* strains** |  |  |
| M145 | Wild-type *S. coelicolor* A3(2) |  |
| WT::pflux-EV | Promoterless *luxCDABE* operon | <sup>31</sup> |
| WT::pflux-hrdBp | *luxCDABE* operon driven by the hrbB sigma factor promoter | <sup>31</sup> |
| WT::pflux-redDp | *luxCDABE* operon driven by the RedD promoter | <sup>31</sup> |
| WT::pflux-eCISp1 | *luxCDABE* operon driven by the eCIS promoter P1 | This study |
| WT::pflux-eCISp2 | *luxCDABE* operon driven by the eCIS promoter P2 | This study |
| WT::pflux-eCISp3 | *luxCDABE* operon driven by the eCIS promoter P3 | This study |
| WT::pflux-eCISp4 | *luxCDABE* operon driven by the eCIS promoter P4 | This study |
| M1152 | Derivative of M1146, Δ*act*, Δ*red*, Δ*cpk*, Δ*cda*, *rpoB* [C1298T] | <sup>22</sup> |
| M145 ΔTS (Δ*sco4253*::*apra*) | eCIS tail sheath gene *sco4253* replaced with apramycin resistance cassette | This study |
| M145 ΔBP (Δ*sco4243*Δ*sco4244*Δ*sco4245*::*apra*) | eCIS baseplate genes *sco4243-45* replaced with apramycin resistance cassette | This study |
| M145 ΔATPase (Δ*sco4259*::*apra*) | eCIS ATPase gene *sco4259* | This study |
|  | replaced with apramycin resistance cassette |  |
| M1152 ΔTS (Δ*sco4253::apra*) | | This study |
| M145::pIJ6902-actR-FLAG | M145 conjugated with pIJ6902 plasmid encoding inducible FLAG-tagged ActR (Cytoplasmic control) | <sup>63</sup> |
| M145::pIJ6902-S4646-FLAG | M145 conjugated with pIJ6902 plasmid encoding inducible FLAG-tagged SecE (Membrane control) | <sup>63</sup> |
| Plasmids |  |  |
| pFlux | Reporter plasmid, Promoterless luxCDABE operon, ApraR | <sup>31</sup> |
| pFlux-eCISp1 | eCIS promoter region P1 (Sco4248-4249 reverse) | This study |
| pFlux-eCISp2 | eCIS promoter region P2 (Sco4248-4249 forward) | This study |
| pFlux-eCISp3 | eCIS promoter region P3 (Sco42453-4254 reverse) | This study |
| pFlux-eCISp4 | eCIS promoter region P4 (Sco42453-4254 forward) | This study |
| pIJ7331 | pBluescript KS (+), aac(3)IV, oriT (RK2) | <sup>55</sup> |
| p15TV-L | T7 expression vector, derivative of pET15b, AmpR, (GenBank ID: EF456736.1) | Novagen |
| p15-SC4245 | 6XHis BW1 expression plasmid | This study |
| p15-SC4253 | 6XHis tail sheath expression plasmid | This study |
| Cosmids |  |  |
| StD8a |  | <sup>57</sup> |
| StD8aΔsco4253::apra | | This study |
| StD8aΔsco4243-45::apra | | This study |
| StD8aΔsco4259::apra | | This study |

**Supplementary Table 4:**
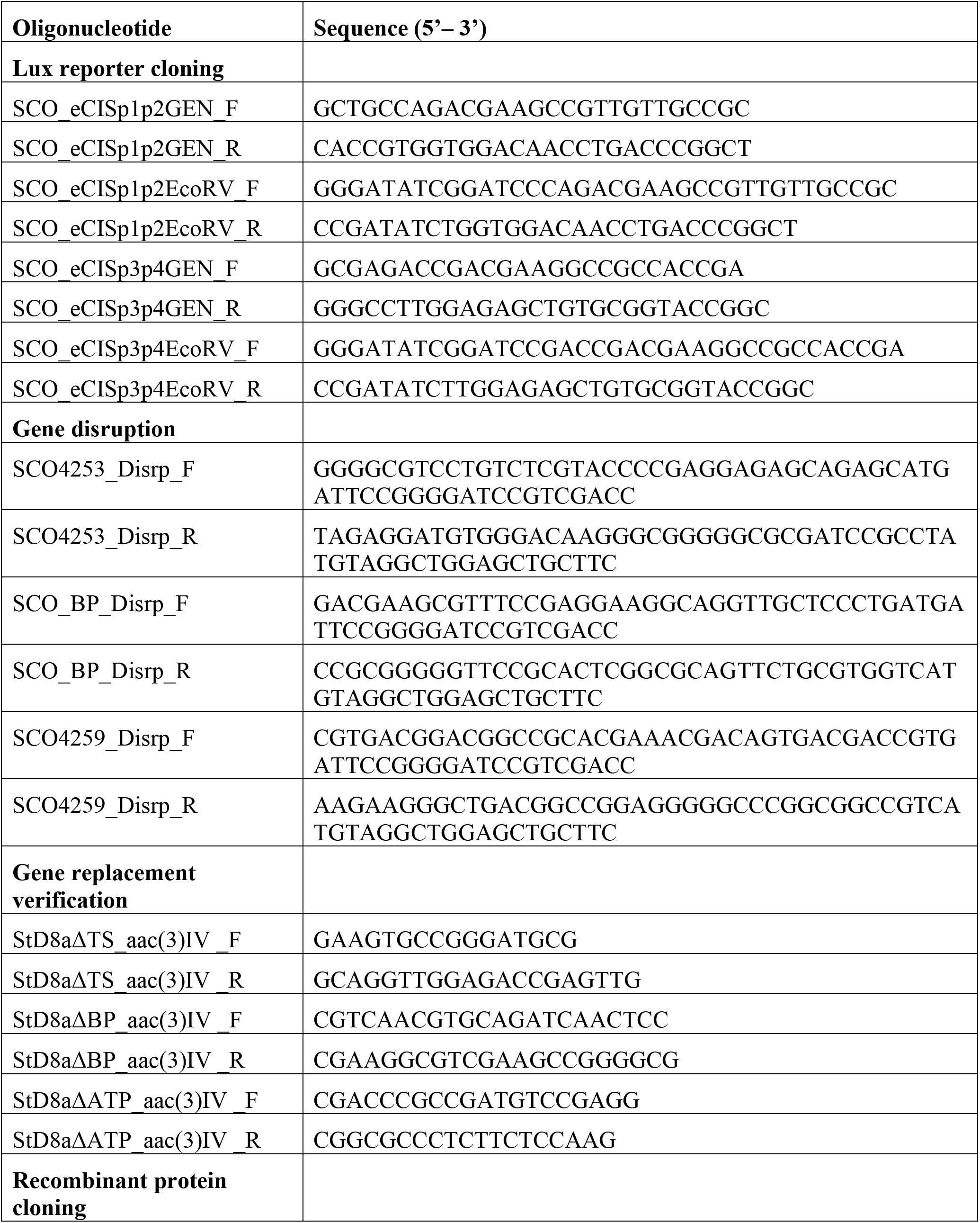
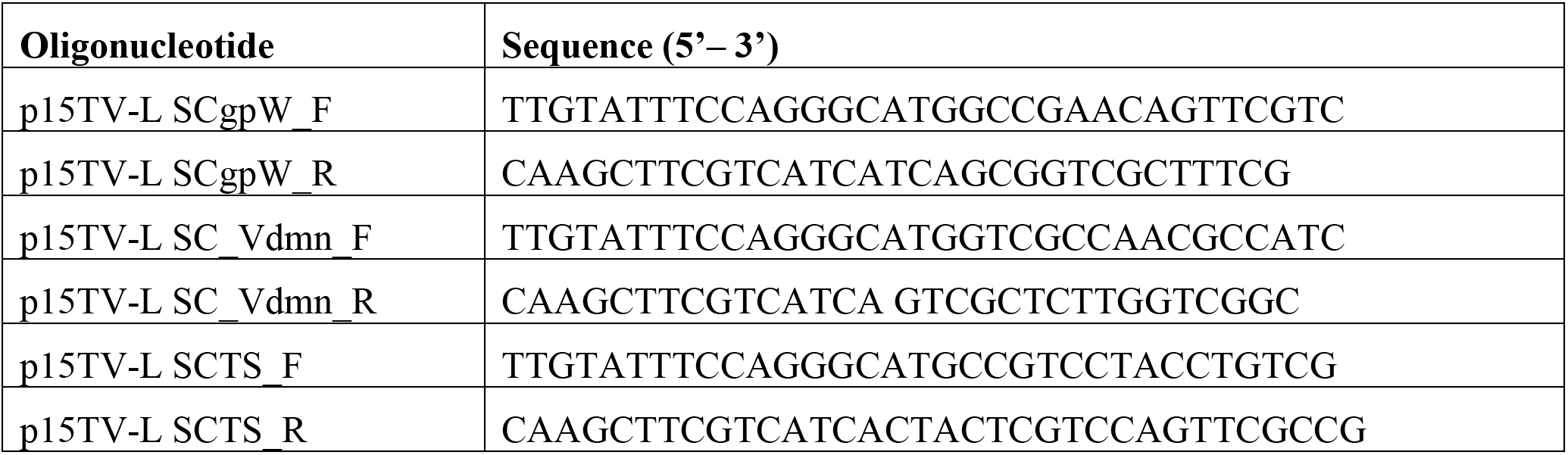
Oligonucleotides used in this study.

